# Refinement and maintenance of receptive field size in primary visual cortex occurs without visual experience in mice

**DOI:** 10.64898/2026.06.27.735014

**Authors:** Pedro Fernández-Aburto, Korey N. Sudana, Sarah L. Pallas

## Abstract

A critical step in visual cortical maturation is refinement of receptive field (RF) size, producing higher acuity vision. This was previously studied using species with well-developed vision (e.g., carnivores, primates), in which visual experience was necessary for refinement but not maintenance of RFs in visual cortex. In contrast, in Syrian hamsters, a crepuscular species with low visual acuity, dark rearing had no effect on RF refinement in juveniles, but RFs re-enlarged in adulthood, resulting in reduced acuity. These inter-species differences raise the question of whether the need for visual experience is primarily related to the phylogenetic position of the species or to its ecological niche. Here we report that dark rearing had no effect on development or maintenance of RF properties of visual cortical neurons in nocturnal mice. Mice with lifelong visual deprivation refined and maintained their RF size over time. Furthermore, the development of stimulus direction tuning was unaffected by dark rearing. In contrast, surround suppression, orientation tuning and the sharpness of direction tuning were abnormal in dark reared mice. These and our previous results from hamsters show that species living in an ecological niche with minimal daylight exposure require little to no visual experience to develop and maintain refined RFs. This study is an important step in developing a better understanding of the role of visual experience in the development of visual processing circuitry and suggests that diurnal mammals may be a better model for human visual cortical development than mice.

**SIGNIFICANCE STATEMENT:** The role of visual experience in development and plasticity of receptive fields in visual cortex was initially studied using animals with well-developed visual systems (carnivores, primates). Mice have been used more recently, primarily because of genetic manipulations. Now that gene alteration is possible in other species, whether mice are the best choice for visual development and plasticity studies needs reevaluation. We assayed the role of visual experience in mouse visual cortical development, focusing on receptive field refinement. We report that visual experience is unnecessary for the refinement of receptive fields in visual cortex, suggesting that mice *(Mus musculus)* may not be the best animal model to demonstrate how visual experience influences the development of visual cortical circuitry in humans.

## INTRODUCTION

It has been argued, based on prior studies of visual deprivation and ocular dominance (OD) plasticity in primary visual cortex (V1) primarily in macaques and domestic cats (see Espinosa and Stryker, 2012, for review), that early visual experience during a critical period is required for development of visual cortical circuits, but not for the maintenance of refined circuitry. For example, visual deprivation in kittens and juvenile monkeys results in reduced responsiveness to light, a lengthening and/or delay of the critical period, and a failure to refine receptive fields (Wiesel and Hubel, 1963, 1965; Blakemore et al., 1978; LeVay et al., 1980; Mower, 1991) . But different receptive field (RF) properties can vary in dependence on early visual experience between different species. In primates and carnivores, sharpening of direction tuning in V1 requires light exposure (as observed in cats (Berman and Daw, 1977; Leventhal and Hirsch, 1980; Sengpiel et al., 1999)) and ferrets (Li et al., 2006; Li et al., 2008; Van Hooser et al., 2014; Roy et al., 2018) . Development of orientation tuning, in contrast, does not require visual experience for development (cats: Sengpiel et al., 1999; (Chapman et al., 1996; Li et al., 2006; Li et al., 2008; Van Hooser et al., 2014; Roy et al., 2018) Furthermore, cats and monkeys do not exhibit ocular dominance plasticity in adulthood (Hubel and Wiesel, 1970; LeVay et al., 1980). Direction tuning can be modified by experience in rats (Fagiolini et al., 1994), but not in mice (*Mus musculus*: Rochefort et al., 2011). In contrast to the carnivores and primates that have been studied, mice exhibit ocular dominance plasticity in adulthood (Hofer et al., 2006). Our previous results in Syrian hamsters revealed that RFs in the superior colliculus (SC) and V1 require visual experience only to maintain refinement in adulthood but not to refine during development (Carrasco et al., 2005; Balmer and Pallas, 2015), opposite from results in cat and monkey studies. These contrasting findings demonstrate that the role of visual experience in the development of receptive field (RF) properties can differ across different species, different visual response properties, and different brain regions.

The maturation of GABAergic lateral inhibition sculpts RF refinement in visual circuits by sharpening spatial tuning and suppressing redundant inputs, and perturbations in this inhibitory drive reliably produce RF enlargement across species (Huang et al., 1999; Berardi et al., 2000; Gandhi et al., 2008; Chen et al., 2014). Dark reared hamsters’ failure to maintain refined visual RFs in adulthood is due to a loss of GABAergic inhibition (Carrasco et al., 2011) which results from decreased BDNF-TrkB signaling (Mudd et al., 2019). The timing of the reduction in GABA levels coincides with the timing of RF enlargement (Battle-Otfinoski & Pallas, in prep.)

Our long-term goal is to understand how these species differences came to exist. The overarching question is whether the role of visual experience in refinement and maturation of RFs in visual cortex is influenced more by ecological niche or by phylogenetic position. In this study our aim was to determine if visual experience is necessary for developmental refinement and/or maintenance of visual cortical RF properties in nocturnal mice. Based on our findings in crepuscular hamsters that DR causes a delayed loss of BDNF/TrkB-mediated GABAergic inhibition (Mudd et al., 2019), the hypothesis tested here is that ecological niche, primarily as related to circadian rhythm (i.e. whether animals are nocturnal, diurnal, or crepuscular), defines the need for visual experience in the maturation of RF size more so than does phylogeny. Although the mouse visual system and the influence of visual experience on development of its visual acuity have been studied extensively (e.g., Gianfranceschi et al., 2003; Prusky and Douglas, 2003; Kang et al., 2013), information concerning RF refinement in mice is limited. Thus, comparative studies about the role of visual experience in the refinement and maintenance of RF size are needed.

This study investigated how visual experience influenced the development of various RF properties in V1 neurons of lab mice *(Mus musculus)*, including RF size, selectivity for orientation and direction of a stimulus, and stimulus size tuning across different postnatal ages. We found unexpectedly that when mice are dark reared from before birth until adulthood, RFs of V1 layer 2/3 neurons are refined to a smaller size as occurs in normally reared mice, and that the refined state is maintained through adulthood. In addition, we observed that direction selectivity did not require visual experience. However, orientation selectivity, sharpness of direction tuning and surround suppression did require light exposure during development. Finally, the normal development of a preference for grating stimuli with a diameter similar to the RF size was negatively affected by dark rearing. Based on these results, we conclude that mice, unlike humans, do not require visual experience to refine and maintain RF size. On the other hand, the refinement of the area surrounding the RF center is affected by early light exposure. Our finding that some RF properties including RF size, which is critical to visual acuity, develop independently of visual experience in mice suggests that it is not an ideal animal model for development of V1 in humans, and that diurnal rodent species are likely to be a better choice for understanding visual cortical development in humans.

## METHODOLOGY

### Animals and rearing conditions

C57BL/6J mice were obtained from Charles River Laboratories and placed in a 12h:12h light/dark cycle, temperature-controlled room (27°C). They were maintained in cages with water and food *ad libitum*. Experiments involved a total of 27 mice reared in normal (NR) or dark (DR) conditions and examined at different postnatal ages: ∼P30 (P25-P45); ∼P60 (P51-P70) and ∼P90 (P90-P120). Normally reared (NR) mice were housed under a 12:12 light:dark cycle. For generating dark reared (DR) mice, pregnant females were placed prior to parturition in dark chambers within a dark room arranged to avoid any light exposure. Husbandry procedures used a far-red light that cannot be seen by mice (Jacobs et al., 1991). Pups were born and reared in dark conditions and were not exposed to light until the recording session began. The NR and DR pups stayed with their mother and siblings until they were weaned at approximately 21 days, after which they were separated by sex. All animal procedures performed in this study were approved by the Institute of Animal Care and Use Committee (IACUC) of the University of Massachusetts-Amherst and followed the Society for Neuroscience recommendations for animal use and the National Research Council (NRC) Guide for the Care and Use of Laboratory Animals.

### Surgery and electrophysiological recording

Mice used for recording were first anesthetized using isoflurane (3% + 400 cc/min O_2_) followed by intraperitoneal injections of urethane (1 g/kg), divided into four doses applied I.P. at 15 min intervals. Chlorprothixene (8 mg/kg), a sedative compound, was injected I.P. 5 min after the first urethane dose. During the period of anesthesia, atropine (0.005 mg/kg) and dexamethasone (1 mg/kg) were administered through subcutaneous and intramuscular injections, respectively. The animals were given a subcutaneous injection of 5% dextrose in Ringer’s solution (0.05 - 0.1 mL) every ∼3 h to maintain hydration. Anesthesia level was evaluated by monitoring reflexes in the tail, toes, ear lobes, and eyelids. A continued absence of reflexes was maintained throughout the experiment by administering small injections of urethane (0.25 g/kg). Body temperature was kept at 37° C with the use of a temperature-controlled heat pad (CWE, Ardmore, PA). Tracheostomy was performed under aseptic conditions, and the trachea was exposed to a continuous supply of oxygen (400 - 450 cc/min) through a tracheostomy tube. The right side of the brain was exposed by taking out an approximately 3 x 3 mm section of the skull above area V1, leaving the dura intact. The caudal edge of the skull piece coincided with the lambda suture. The electrode was adjusted by age-appropriate distances from the starting point (lambda) to a position from -0.5 – to 1.0 mm in the rostrocaudal direction and from 2.5 - to 4.0 mm in the mediolateral direction. Tungsten electrodes with a resistance of 1.5 MOhm (FHC, Bowdoin, Maine) were used for recordings at depths ranging from 100 to 300 µm in 5-13 different locations within each subject, with a minimum distance of 50 µm between each location. To ensure proper placement of the electrode in V1, various locations were tested prior to recording. The visual field locations used in our analysis were located within 10° and 45° in azimuth, which includes the binocular portion of the visual field (<30°), and part of its monocular portion (>30°) (Dräger and Olsen, 1980), and between -10° and +30° in elevation similar to those chosen in previous mouse studies (Drager, 1975; Wagor et al., 1980; Hübener, 2003).

### Stimuli

All stimuli used in this study were generated using customized MATLAB scripts. These were presented on a screen placed 25 cm from the animal’s left eye and adjusted to ensure that the receptive fields of the neurons recorded were near the center of the screen. The stimuli were adjusted to align with the RF center before each presentation.

#### Flashing square

A computer screen of 1920 x 1080 pixels size was positioned in front of the left eye and adjusted both to align the RFs with the grid and to minimize stimulus eccentricity (distance of visual stimulus from the RF center) that could affect visual processing. To assess the RF size and response properties of neurons in layer 2/3 of V1, a small white square (60 pixels, 3.4° size) was presented over a black background for 0.15 s at different locations, positioned randomly on a 900 x 900 pixel grid (52° x 52°) with a 0.05 s interstimulus interval (ISI). Each stimulus was repeated 20 times at each location in the grid.

#### Circular sinusoidal drifting grating

To assess the responses of neurons in layer 2/3 of V1 to the orientation and direction of a stimulus, a circular stimulus containing a sinusoidal grating moving in 8 different directions (0-315°, with 45° steps) was shown for 4 s and repeated 10 times for each direction, with a 3 s ISI. The background screen was gray. The size, spatial frequency, and temporal frequency of gratings were kept constant at a radius of 15.2°, 0.04 cycles per degree (cdp), and 1.5 Hz, respectively, for every trial. The preferred direction, which is the direction angle of grating movement at which neurons recorded showed the maximum firing rate was determined at the end of each presentation and then utilized in the size tuning assays.

#### Filled and ring shaped drifting gratings

Circular stimuli containing sinusoidal gratings of different sizes, either a filled circle (radius: 1.7°, 3.5°, 4.3°, 8.6°, 13.8°, 17.3°, 23° and 28.8°) or a ring shaped annulus (external radius: 31.1°; internal radius: 1.7°, 3.5°, 4.3°, 8.6°, 13.8°, 17.3°, 23° and 28.8°) moving in the preferred direction (estimated from data in the orientation/direction selectivity assay) were randomly presented 5 times for 4 s, with 3 s ISI. The background screen was gray. The spatial and temporal frequency of the gratings were kept constant at 0.04 cpd and 1.5 Hz, respectively, for every trial (Popović et al., 2018).

### Data collection and spike sorting

Responses captured from multiunit recordings were sampled at 25 kHz and collected using the Spike2 program (CED, Milton, U.K.). This raw, digitized data was filtered using a band pass filter (Finite impulse response, 700-7000 Hz). To follow the same single unit across the different stimuli used, the templates for the spike waveforms obtained in the first assay (RF size) at each site recorded were retained and used for matching against waveforms in the following assays (orientation/direction selectivity and size tuning). The threshold used for the sorting of single units was obtained from the voltage variance across the time of the stimulus presentation, with the threshold used corresponding to 4 times the calculated root mean square (RMS) of voltage for the duration of the stimulus presentation. Spikes chosen for inclusion in the data set exceeded 4xRMS and were assessed over a time window of approximately 1.5 ms. The single unit data from normal and DR mice were compared after k-means clustering.

### Data analysis

The analysis of responses to each stimulus was performed using customized scripts created in MATLAB. Most of the functions used to fit the data were obtained from Git Hub repository of the van Hooser laboratory (https://github.com/VH-Lab). In order to collect data from layer 2/3, single units located 100 - 300 µm deep were selected for the analysis (**Fig. 1**).

**Figure 1.**
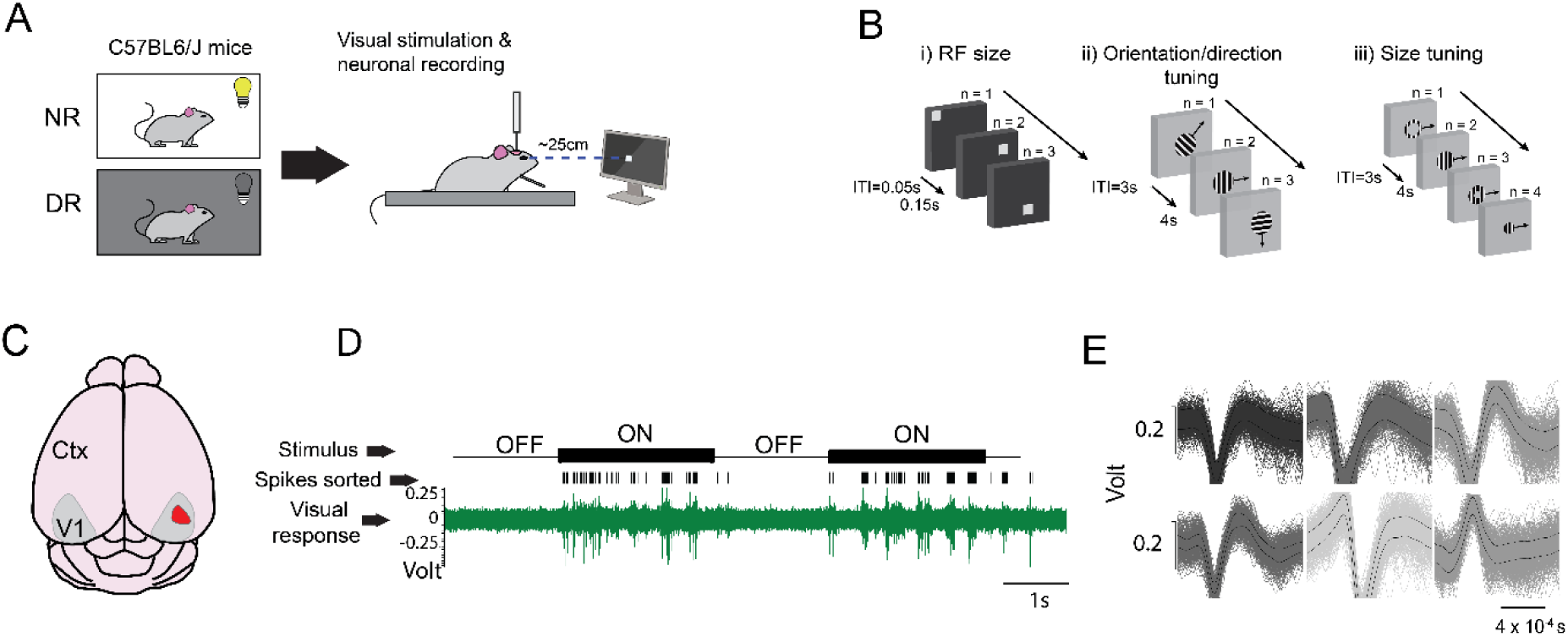
Experimental procedure for the isolation of V1 neurons. **(A)** Schematic of the recording procedure. V1 layer 2/3 neurons were recorded in anesthetized NR and DR mice of different ages. **(B)** Schematics of the different stimuli presented for RF size, orientation/direction tuning and size tuning measurements. **(C)** Schematic of the dorsal view of mouse brain showing the V1 areas (gray) in both hemispheres of cerebral cortex (Ctx). Red area marks the location of recording sites within V1. **D)** Example of an *in vivo*, extracellular recording. Responses of V1 neurons (green) to drifting gratings presented at different orientations were obtained using extracellular recordings. Spikes were sorted and the responses of single units were isolated (black tick marks). **(E)** Examples of isolated V1 single unit responses after sorting and clustering analysis.

#### RF size and shape

The mean firing rate of each neuron for each location on the grid was calculated by averaging its responses to each stimulus type across 20 repetitions. The RF sizes included in the analysis were obtained from single units that had a firing rate equal to or over 1 spike/s after subtraction of spontaneous activity. The spontaneous firing rate was measured 1 s before any stimulus presentation. A 2-D Gaussian function was fitted for each single unit response to obtain the RF size. RF size corresponds to the ellipse function that circles the 2D-Gaussian function at half of its amplitude. RF size contour was defined by the ellipse formula:

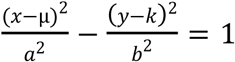

µ and *k* correspond to the mean values describing the 2D-Gaussian function*. a* and *b* correspond to the half-length of the major and minor axis of the ellipse, respectively. RF half-length corresponds to the half-length of the major axis of each RF. RF shape was evaluated by calculating the index (major half-length axis/minor half-length axis), with an index of 1.0 being circular (Márquez et al., 2025). The mean responses were normalized before the curve was fitted by setting the peak of the mean response to 1 (as in Mudd et al., 2019) to account for differences in response strength across single units. Goodness of fit was evaluated by calculating the R^2^ value and including only those single units showing a R^2^ equal to or higher than 0.5.

#### Orientation and direction selectivity

The orientation and direction tuning curves were fitted using the mean response at each angle of grating motion after subtraction of the spontaneous firing rate that was obtained 0.5 s before any stimulus presentation. Tuning curves were designed as two Gaussians of the same width in a circular space that were forced to peak 180° apart (Carandini and Ferster, 2000; Mazurek et al., 2014; Popović et al., 2018).

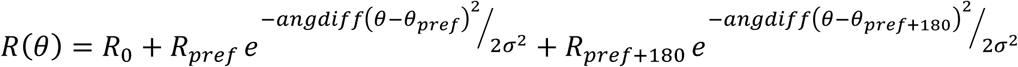

The goodness of fit of the function to the data was evaluated by calculating the R^2^ value and including only those single units showing a R^2^ value equal to or higher than 0.8. The absolute angular difference in circular space is represented as angdiff(*θ*). Orientation selectivity was quantified as the Orientation Index (OI), which is the normalized difference between the sum of responses to the preferred orientation vs. the sum of responses to the opposite orientation as follows: OI= (R_pref_ + R_pref +180_) - (R_pref+90_ + R_pref +270_) / (R_pref_ + R_pref +180_). Direction selectivity was determined as the normalized difference between the response to the preferred direction vs. that of the opposite direction (Direction Index (DI) = (R_pref_ - R_pref +180_) / (R_pref_)). From the total of V1 neurons used for orientation and direction selectivity analyses, the percentage of single units showing high selectivity for orientation or direction (OI and DI > 0.5, respectively) was calculated. We also calculated the sharpness of the direction tuning curve for each single unit, which corresponds to the half width of the curve at half of the maximum amplitude of firing rate.

#### Linearity

The linearity of single unit responses was determined by analyzing the responses to drifting gratings moving in the preferred direction. This was done by dividing the response during the first two seconds of each grating presentation into 0.1 s intervals and subtracting the spontaneous firing rate. Then, a discrete Fourier transform was used to calculate the ratio of response at the preferred drift frequency (F1) to the average response (F0) (median < 0.5 spikes/s, Niell and Stryker, 2008). Neurons showing a F1/F0 ratio higher than 1 were classified as simple neurons and those showing a F1/F0 ratio lower than 1 were classified as complex neurons.

#### Size tuning

To quantify the V1 neurons’ responses to stimulation of the surround, a function that included a Gaussian center component (the response at the RF center) was fitted, then modulated by a circular surround component (filled and ring-shape gratings of different sizes). Filled gratings of 8 different diameters (1.75° to ∼30°) were presented, and the same values were used as internal diameters for ring shaped gratings, similar to the protocol used in Popović et al. (2018). Each stimulus was presented in random order 5 times for 4 s in each session. The evoked firing rate was obtained after subtraction of the spontaneous firing rate. Spontaneous firing rate corresponded to the average baseline response obtained during 0.5 s before the presentation of each stimulus. Neurons exhibiting a Gaussian fit R^2^ > 0.7 were used for the analysis.

To quantify size tuning, we used a fit function that incorporated a Gaussian center component modulated by a circular surround component (Popović et al., 2018). Responses were fitted to a product of two functions that represented the response to the stimulus at the center of the receptive field to a stimulus *S*(*x,y*) (*R_center_*) and the modulation of the center response (*R_mod_*) to stimuli with a different radius (*r*), respectively, as follows:

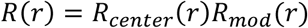

The response of each single unit was fitted using a Gaussian function.

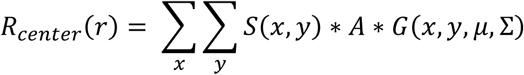

Here *r* is the stimulus radius, *A* is the maximum amplitude of the response (in spikes/s), μ is the center position of the stimulus, which coincided with the center position of the RF, and Σ is the covariance matrix, which indicates how sharp or flat the fitted Gaussian (*G*) is, as well as its orientation along both axes. The Gaussian function was constrained to a circular shape due to the nature of the stimulus used (circular grating patches) by using the following definition for the covariance matrix.

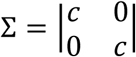

The value *c* corresponds to the standard deviation at half of the maximum response of the Gaussian function fitted for each V1 neuron. RF size corresponds to the circle at half-height of the *G* function. The modulating function *R_mod_*takes values between 0 and 2 and is proportional to the overlap of the stimulus and a circle of radius as *R_max_stim_ as* follows:

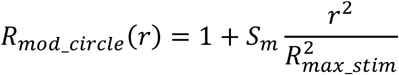

When the stimulus was a ring (circle with central aperture), the modulating response was as follows:

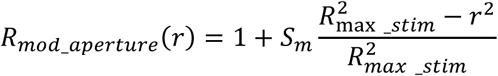

We quantified size tuning using measuring the size modulation parameter (*S_m_*), which represents the degree and direction of modulation of the response at the RF center by surround stimulation. Positive *Sm* values indicate length summation, whereas negative values indicate surround suppression. We also quantified the modulation of the responses to stimuli at the center of each RF by surround stimulation as the surround suppression ratio (SR) (Wang et al., 2010). This analysis only considers the V1 neuron responses to filled gratings of different radii (1.75° to ∼30°). SR corresponds to the ratio of the response to the largest stimulus (R_largest_size_) over the response to the stimulus giving the maximum response (R_preferred_size_), as the following:

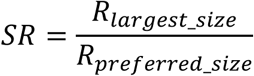

If the stimulus producing the maximum response is the largest stimulus, then SR = 1 and little to no modulation of the neuronal response is observed. If instead a smaller stimulus is producing the maximum response, and thus SR < 1, there is a suppressive effect on the neuronal response. Finally, we evaluated the relationship between the size of the stimulus evoking the maximum response and the RF size of V1 neurons. This was done by calculating the distribution of the differences between RF center size and the size of the stimulus that causes the maximum response.

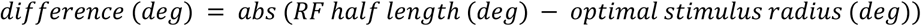

with *abs* as the absolute value.

### Statistical analysis

All data were tested for normality using the Kolmogorov-Smirnov function. Parametric (*t*-test) and non-parametric (Mann-Whitney *U* test) statistical tests were used to compare between rearing groups at each age evaluated. Kruskal-Wallis or 1-way ANOVA and post hoc tests (e.g. Holm-Sidak and Dunn’s test for parametric and non-parametric analysis, respectively) were used to compare across ages in both rearing groups. A *p* ≤ 0.05 was considered statistically significant.

## RESULTS

It is generally thought, based on the work of Hubel and Wiesel (1962), that early visual experience is necessary for the appropriate maturation of most receptive field properties. However, our previous work on hamsters (Carrasco et al., 2005; Carrasco and Pallas, 2006; Carrasco et al., 2011; Balmer and Pallas, 2015; Mudd et al., 2019) suggests that its necessity may vary across species belonging to different phylogenetic groups and/or having different visual habits. A detailed description of the role of visual experience on the development of visual properties across species would help to clarify the issue. In this study, we evaluated the effect of visual experience on the development and maintenance of receptive field properties in visual cortex of nocturnal mice *(Mus musculus)*. Starting at ∼P30 and continuing until adulthood, the responses of V1 neurons in layer 2/3 to different stimuli were obtained from normally reared mice and mice reared in the dark from before birth until recording. For each age and rearing group, the RF size and the tuning for stimulus orientation, direction and size were examined. Based on our analysis of the linearity of V1 neurons’ responses (shown below), neurons across ages and rearing conditions exhibited non-linear responses to oriented, grating stimulus.

### Rearing-dependent differences in spontaneous activity emerge at puberty but are absent by adulthood

Before examining the effect of visual experience on the development of neuronal activity in V1 layer 2/3 neurons, we evaluated the spontaneous activity rates of single units across ages and between normal and dark rearing conditions. Spontaneous activity rates in the adult NR mice in our study (median ranging 0-1 spikes/s) (**Table 1**) were within the range of rates previously reported by others (median 0.5 spikes/s, Niell and Stryker, 2008; median = 2.7 spikes/s, Gao et al., 2010, Drager, 1975). At ∼P30 there was no significant difference in spontaneous activity across rearing conditions (Mann-Whitney *U* test = 581, *p* = 0.3) (**Table 1**)). At ∼P60, a significantly higher spontaneous firing rate was observed in DR mice than in NR mice (Mann-Whitney *U* test = 901, *p* = 0.014) (**Fig. 2A**). However, by ∼P90 there was no significant difference between NR and DR mice (Mann-Whitney *U* test = 838, *p* = 0.2). Thus, spontaneous activity levels in V1 neurons differ between pubertal NR and DR mice, but not in adults that have or have not received visual experience.

**Figure 2.**
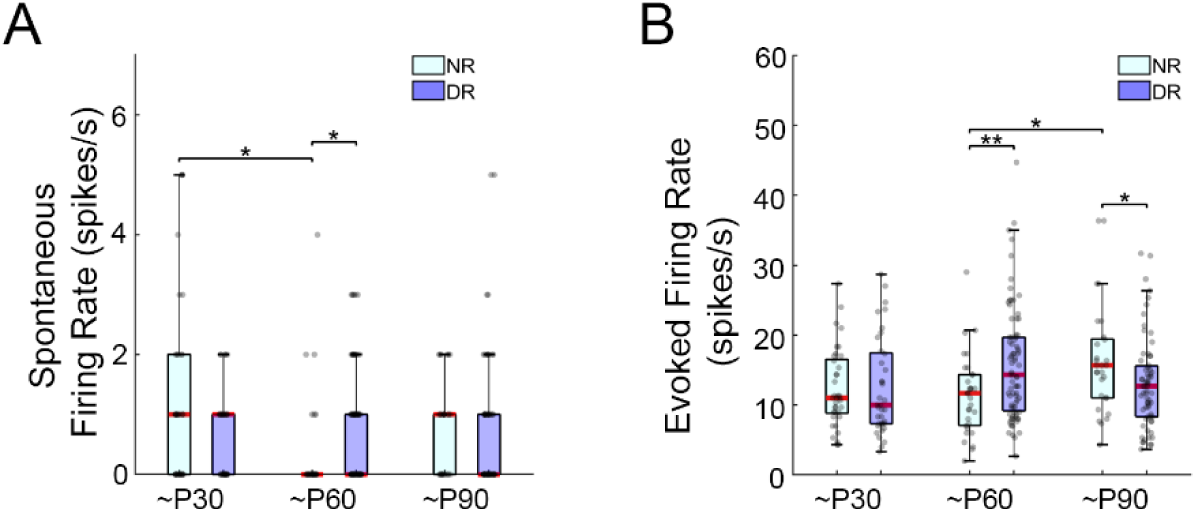
Response properties of NR and DR mouse V1 neurons in layer 2/3 across postnatal development. Layer 2/3 neurons in mouse V1 exhibited spontaneous activity across ages and rearing conditions. Evoked response levels, but not spontaneous activity levels, were affected by visual experience. **(A)** Mean spontaneous firing rate and **(B)** mean evoked firing rate of single units during stimulation of the RF center. Medians are shown in the boxplots (red lines) along with confidence intervals for single units with 25th and 75th percentiles. Whiskers correspond to minimum and maximum values within a 1.5x interquartile range. *p<0.05, **p<0.01, ***p<0.001

**Table 1.**
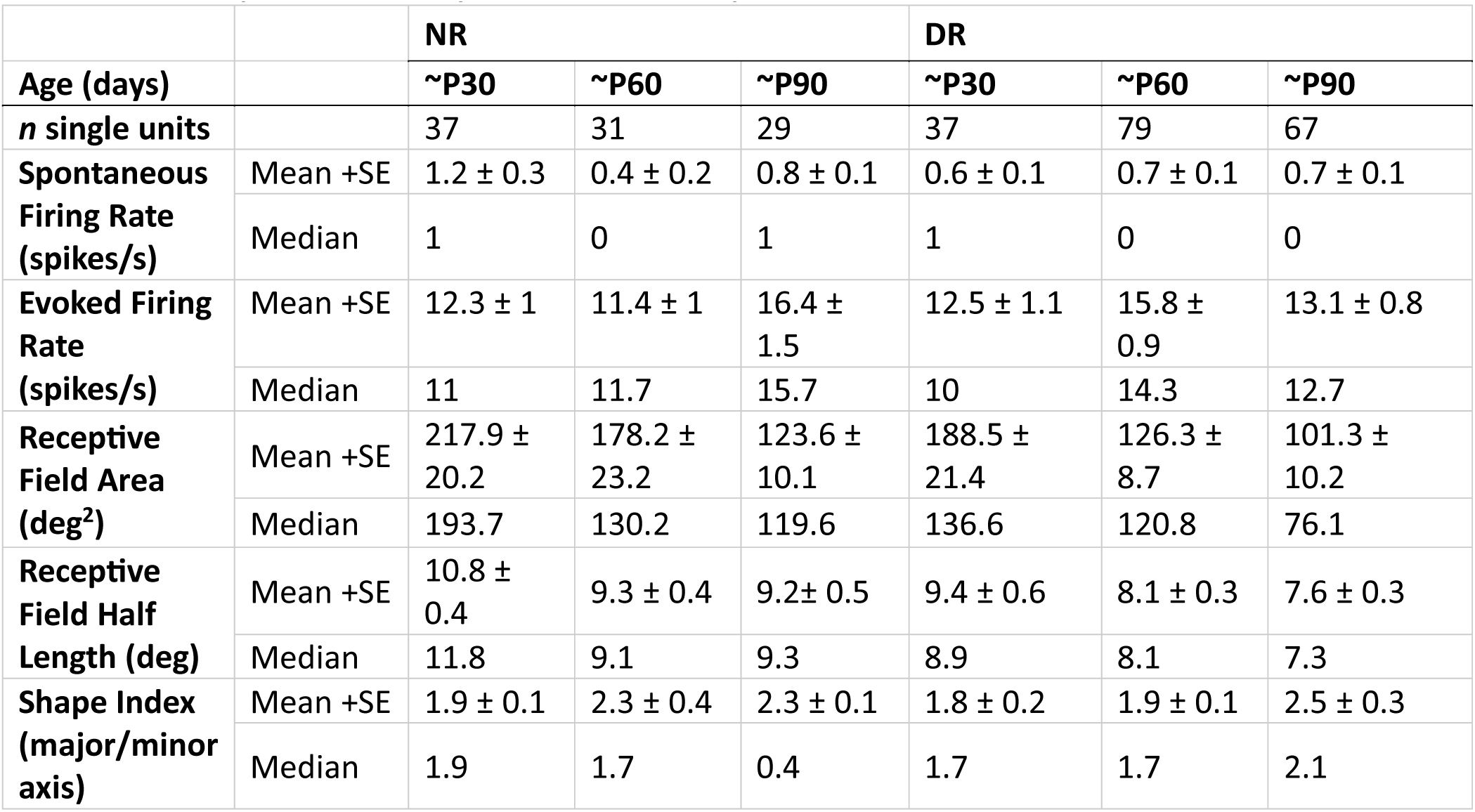
Comparison of receptive fields and response levels between NR and DR mice.

|  |  | NR |  |  | DR |  |  |
| --- | --- | --- | --- | --- | --- | --- | --- |
| Age (days) |  | ~P30 | ~P60 | ~P90 | ~P30 | ~P60 | ~P90 |
| ***n* single units** |  | 37 | 31 | 29 | 37 | 79 | 67 |
| **Spontaneous Firing Rate (spikes/s)** | Mean +SE | 1.2 ± 0.3 | 0.4 ± 0.2 | 0.8 ± 0.1 | 0.6 ± 0.1 | 0.7 ± 0.1 | 0.7 ± 0.1 |
|  | Median | 1 | 0 | 1 | 1 | 0 | 0 |
| **Evoked Firing Rate (spikes/s)** | Mean +SE | 12.3 ± 1 | 11.4 ± 1 | 16.4 ± 1.5 | 12.5 ± 1.1 | 15.8 ± 0.9 | 13.1 ± 0.8 |
|  | Median | 11 | 11.7 | 15.7 | 10 | 14.3 | 12.7 |
| **Receptive Field Area (deg<sup>2</sup>)** | Mean +SE | 217.9 ± 20.2 | 178.2 ± 23.2 | 123.6 ± 10.1 | 188.5 ± 21.4 | 126.3 ± 8.7 | 101.3 ± 10.2 |
|  | Median | 193.7 | 130.2 | 119.6 | 136.6 | 120.8 | 76.1 |
| **Receptive Field Half Length (deg)** | Mean +SE | 10.8 ± 0.4 | 9.3 ± 0.4 | 9.2 ± 0.5 | 9.4 ± 0.6 | 8.1 ± 0.3 | 7.6 ± 0.3 |
|  | Median | 11.8 | 9.1 | 9.3 | 8.9 | 8.1 | 7.3 |
| **Shape Index (major/minor axis)** | Mean +SE | 1.9 ± 0.1 | 2.3 ± 0.4 | 2.3 ± 0.1 | 1.8 ± 0.2 | 1.9 ± 0.1 | 2.5 ± 0.3 |
|  | Median | 1.9 | 1.7 | 0.4 | 1.7 | 1.7 | 2.1 |

### V1 neurons in adults exhibited stronger evoked responses in NR mice than in DR mice

The effect of visual experience on the development of stimulus-triggered activity was analyzed by quantifying the neuronal response of V1 layer 2/3 neurons to visual stimuli (evoked response) across age groups and rearing conditions (see **Table 1**). At ∼P30, evoked firing rates in DR animals did not show significant differences from those in NR animals (Mann-Whitney *U* test = 578, *p* = 0.21) (**Fig. 2B**). However, in ∼P60 DR mice V1 neurons exhibited higher mean responses than V1 neurons in ∼P60 NR mice (Mann-Whitney *U* test = 839, *p* = 0.01). The evoked response rates of NR V1 neurons at ∼P90 were higher than those of DR V1 neurons (Mann-Whitney *U* test = 698, *p* = 0.03). From ∼P60 to ∼P90, a significant increase in evoked response rates in NR V1 neurons compared to response levels at ∼P30 was observed (Kruskal-Wallis 1-way ANOVA on ranks, H(2) = 8.2, *p* = 0.02, Dunn’s post hoc test, *p* < 0.05). No significant change was found between ∼P60 and ∼P90 DR subjects (Kruskal-Wallis 1-way ANOVA on ranks, H(2) = 6.7, *p* = 0.04, Dunn’s post hoc test, *p* > 0.05). Overall, light rearing caused stronger evoked activity in V1 neurons than in neurons from mice reared in prolonged darkness.

### Receptive field size of layer 2/3 neurons in V1 refined independent of visual experience

To determine whether visual experience is required for RF refinement, the normalized responses obtained from stimuli presented to the RF areas of each single unit (after subtraction of spontaneous activity) were fitted to 2-D Gaussian curves (**Fig. 3A**). In total, 280 single units that exhibited well fitted curves (R^2^ > 0.5) with clearly discrete RFs were used in this analysis, with 97 single units from NR mice and 183 single units from DR mice (**Table 1**). Neurons with low discharge rates to visual stimulation (< 1 spike/s; 133 single units in total, 56 single units from NR mice and 77 single units from DR mice) were excluded from the analysis.

**Figure 3.**
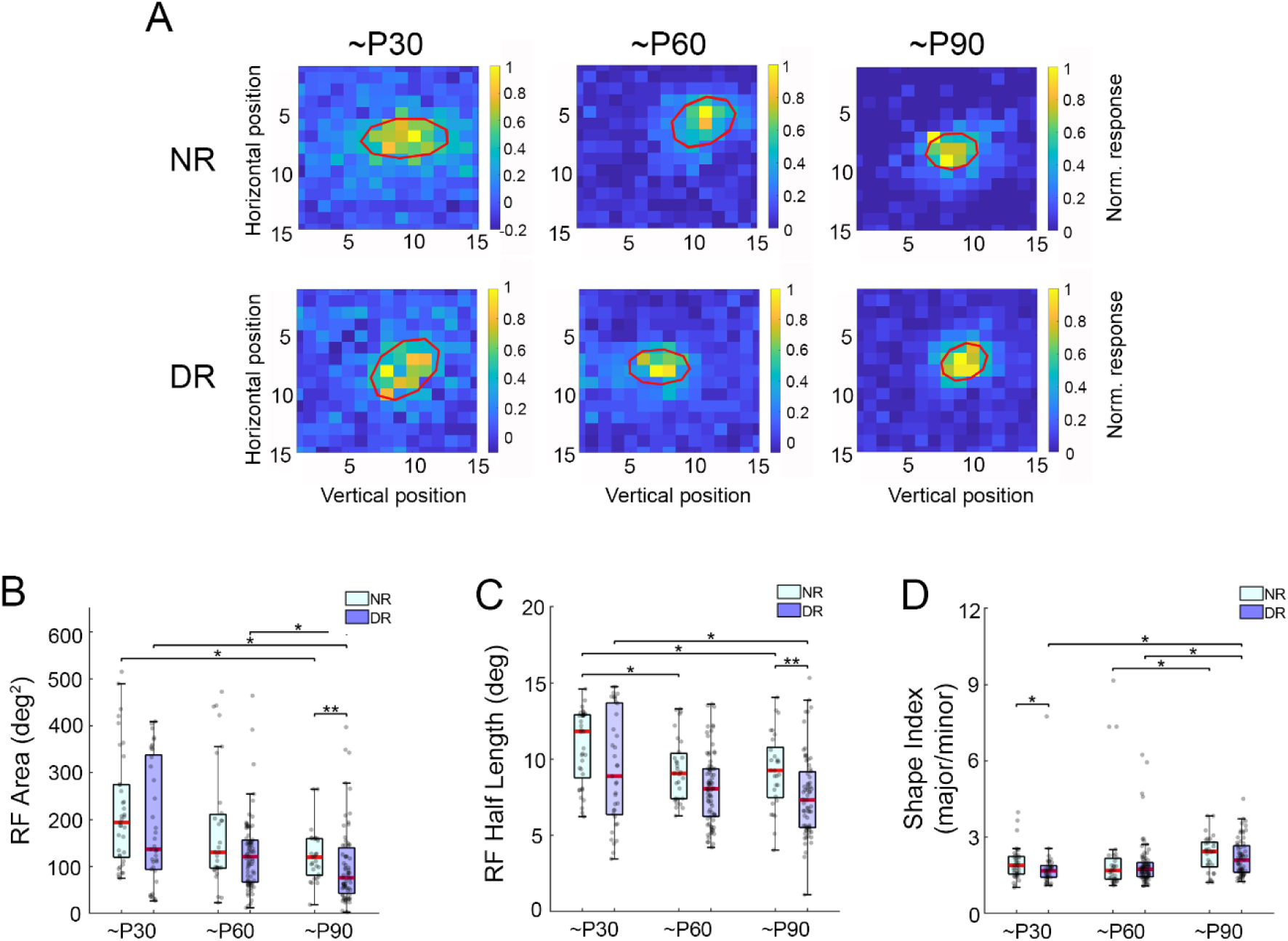
V1 neurons in normally-reared and dark reared mice exhibited similar decreases in their RF size and shape with age. **(A)** Heatmaps show the normalized mean evoked firing rate ± S.E.M at each stimulus location on the screen. RF areas (red ellipse) were calculated by fitting a 2D-Gaussian. Colors indicate relative spikes/s. **(B-D)** Size and shape of RFs from V1 neurons **(B)** Medians in the boxplot (red lines) along with confidence intervals for single unit RF areas with 25th and 75th percentiles. Whiskers correspond to minimum and maximum values within a 1.5x interquartile range. **(C)** Boxplot showing single unit RF half lengths for NR and DR groups across ages.(**D**) Boxplot showing single unit shape index for NR and DR groups across ages. *p<0.05.

Results from other species (rats (Fagiolini et al., 1994), hamsters (Carrasco et al., 2005; Balmer and Pallas, 2015), ferrets (Popović et al., 2018; Fernandez-Aburto et al., 2024), cats (Blakemore and Van Sluyters, 1975; Hirsch, 1985), and monkeys (Jacobs and Blakemore, 1988; Kiorpes, 2015)() predicted that RF refinement would be negatively affected by rearing in darkness, but that was not the case in mice. The RF areas of V1 neurons in ∼P30 NR and DR animals did not show significant differences (NR vs. DR: Mann-Whitney *U* test = 568, *p* = 0.21) (**Fig. 3B, C**, **Table 1**). The same was true in the ∼P60 NR and DR groups (NR vs DR: Mann-Whitney *U* test = 984, *p* = 0.1). However, at ∼P90 RF areas in NR mice were significantly larger than RF areas in ∼P90 DR mice (NR vs DR: Mann-Whitney *U* test = 635, *p* = 0.007). Thus, the RF sizes in mature adults were smaller in mice raised in prolonged darkness than in mice raised in normal conditions.

Across ages, there was a gradual refinement of RF areas in both NR and DR mice. A reduction in RF area between ∼P30 and ∼P90 was observed in NR mice (NR: Kruskal-Wallis 1-way ANOVA on ranks, H(2) = 11.4, *p* = 0.003, Dunn’s post hoc test, *p* < 0.05) (**Figs. 3B, C, Table 1**). As seen in NR mice, a refinement in RF area across ages was observed in DR mice. Like NR mice, DR mice exhibited a significant reduction in RF size between ∼P30 and ∼P90 and between ∼P60 and ∼P90 (DR: Kruskal-Wallis 1-way ANOVA on ranks, H(2) = 15.2, *p* < 0.001, Dunn’s post hoc test, *p* < 0.05). These results suggest that the refinement of layer 2/3 neuron RFs in V1 occurs throughout development in mice, independent of visual experience.

RF half-length was defined as half of the major axis of each RF in NR and DR mice. At ∼P60 and ∼P90 the RF half-length was greater in NR mice than in DR mice (∼P60: NR vs DR: Mann-Whitney *U* test = 857, *p* = 0.02; ∼P90: NR vs DR: *t*-test = 2.8, *p* = 0.007) (**Fig. 3C**, **Table 1**). In contrast, no significant differences in the RF half-length between NR and DR were present at ∼P30 (NR vs DR: Mann-Whitney *U* test = 547, *p* = 0.1). Thus, the resulting half-length RFs in adulthood were reduced in size to a greater extent in mice raised in prolonged darkness than mice raised in 12 light:12 dark conditions.

The RF half-length also decreased across ages in NR and DR mice, with a significant reduction from ∼P30 to ∼P90 (1-way ANOVA, F = 5.6, *p* = 0.005, Holm-Sidak post hoc test, *p* = 0.014) and from ∼P60 to ∼P90 in NR mice (1-way ANOVA, F = 5.6, *p* = 0.005, Holm- Sidak post hoc test, *p* = 0.012) and from ∼P30 to ∼P90 in DR mice (Kruskal-Wallis 1-way ANOVA on ranks H(2) = 6.4, *p* < 0.04, Dunn’s post hoc test, *p* < 0.05) (**Fig. 3C**, **Table 1**). Overall, these results suggest that the refinement of RFs in V1 layer 2/3 neurons occurs independent of visual experience, unlike in cats, in which visual experience is needed during development (Hubel and Wiesel, 1962), and also unlike in hamsters, in which visual experience is necessary in adulthood (Balmer and Pallas, 2015).

### V1 neurons exhibited elongated V1 RFs across ages and rearing conditions

We also investigated the effect of visual experience on the RF shape of layer 2/3 neurons in V1. As in other mammalian species studied, RFs are elongated in V1 of adult mice (Hubel and Wiesel, 1962; Chapman et al., 1999; Gao et al., 2010) . Across ages, RF shape was evaluated by calculating the index (major axis/minor axis), with an index of 1.0 being circular and higher values meaning an increase in length. Neurons exhibited elliptical RF shapes across all ages and light conditions evaluated (**Table 1**). NR mice had more elongated, elliptical RFs in V1 than DR mice at ∼P30 (Mann-Whitney *U* test = 497, *p* = 0.043). However, this difference did not persist across later ages, either at ∼P60 (Mann-Whitney *U* test = 1135, *p* = 0.1) or at ∼P90 (Mann-Whitney *U* test = 857, *p* = 0.4) (**Fig. 3D)**. Instead, a significant increase in the RF shape index was observed from ∼P60 to ∼P90 for both NR RFs (Kruskal-Wallis 1-way ANOVA on ranks H(2) = 9.9, *p* = 0.007, Dunn’s post hoc test, *p* < 0.05) and DR RFs (Kruskal-Wallis 1-way ANOVA on ranks H(2) = 16.4, *p* < 0.001, Dunn’s post hoc test, *p* < 0.05). Thus, the elliptical shape of RFs in V1 layer 2/3 neurons develops independent of visual experience.

### Visual experience was necessary for development of orientation but not direction selectivity

Niell & Stryker (2008) found that layer 2/3 neurons in V1 of mice ranging from 2 - 6 months of age exhibit a strong preference for stimulus orientation but a weaker preference for direction of motion of either bars or gratings. In this study we have assessed the role of visual experience in the establishment of selectivity for orientation and direction. Selectivity for orientation and direction was assessed across ages and rearing groups with a circular stimulus containing a sinusoidal grating moved in 8 different directions (see Methods). In all groups, individual neurons with well-fitted tuning curves for orientation and/or direction were present (**Fig. 4A**).

**Figure 4.**
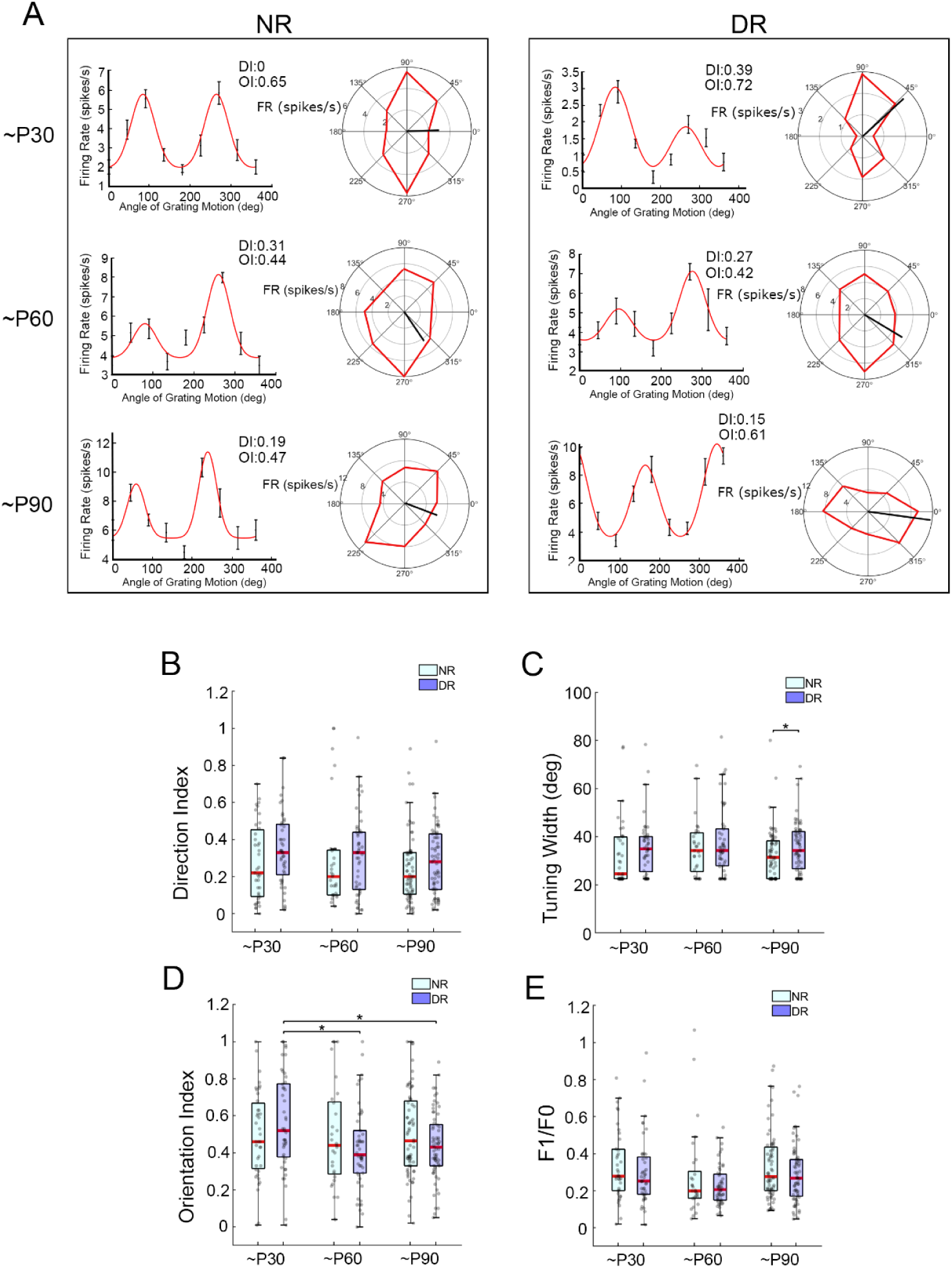
Visual experience was necessary for development of orientation selectivity but not direction selectivity. **(A)** Examples of single unit tuning curves in NR (left) and DR (right) mice at ∼P30, ∼P60, and ∼P90 showing mean evoked firing rate (FR) ± S.E.M to gratings presented at 8 different angles. Linear (left) and polar (right) plots of direction tuning show the response at each angle of grating motion in spikes per second (red line). Bold black lines indicate the vector derived from the mean of responses to individual directions. **(B-E)** Boxplots for **(B)** direction index, **(C)** direction tuning width, defined as the half width at the half maximum response from the direction tuning curve, **(D)** orientation index, and **(E)** F1/F0 ratio for NR and DR groups across ages. Each box shows the median (red lines) and 25th and 75th percentiles. Whiskers correspond to minimum and maximum values within a 1.5x interquartile range. *p<0.05.

#### Direction selectivity

Direction selectivity was observed in under 20% of neurons in both NR and DR mice across ages. Direction selectivity was calculated as direction index (DI), which is the normalized difference between the response to the preferred direction vs. that of the opposite direction. From the layer 2/3 V1 neurons assessed for direction selectivity in NR mice, the percentage of single units with DI > 0.5 was 18% at ∼P30 (7/39 neurons), 17% at ∼P60 (5/29 neurons) and 7% at ∼P90 (5/68 neurons), similar to what has been reported previously (Rochefort et al., 2011). In DR mice, the percentage of single units with DI > 0.5 was 20% at ∼P30 (9/45 neurons), 16% at ∼P60 (8/50 neurons) and 9% at ∼P90 (6/65 neurons). Across all V1 neurons analyzed, DI values were not significantly different between DR and NR groups for any of the three age groups (∼P30: Mann-Whitney *U* test = 702, *p =* 0.1; ∼P60: Mann-Whitney *U* test = 613, *p =* 0.3; ∼P90: Mann-Whitney *U* test = 1905, *p* = 0.2)(**Fig. 4B**, **Table 2**). Thus, the degree of preference for stimulus movement direction was not affected by visual experience.

**Table 2.**
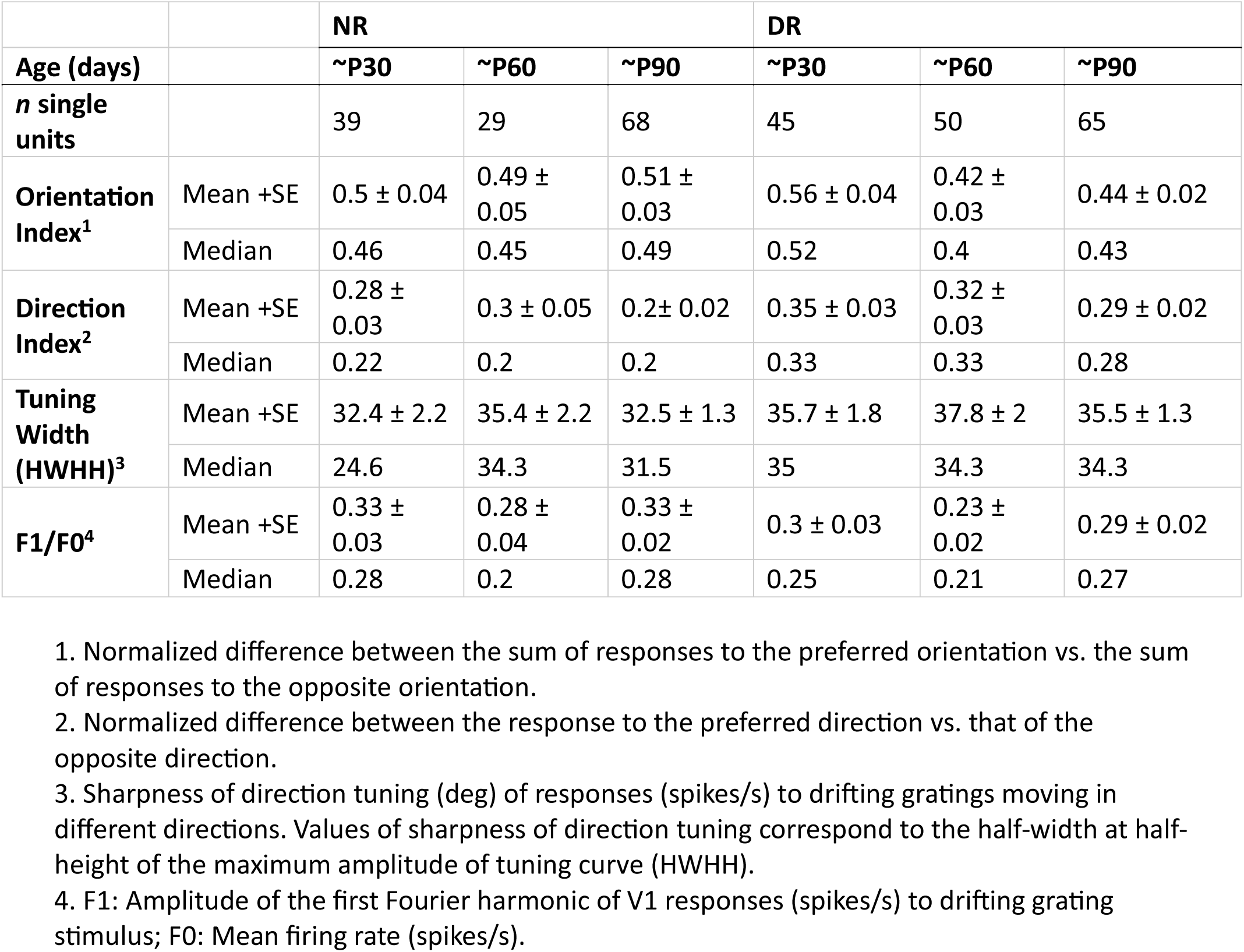
RF properties assayed with drifting gratings.

|  |  | NR |  |  | DR |  |  |
| --- | --- | --- | --- | --- | --- | --- | --- |
| Age (days) |  | ~P30 | ~P60 | ~P90 | ~P30 | ~P60 | ~P90 |
| ***n* single units** |  | 39 | 29 | 68 | 45 | 50 | 65 |
| **Orientation Index<sup>1</sup>** | Mean +SE | 0.5 ± 0.04 | 0.49 ± 0.05 | 0.51 ± 0.03 | 0.56 ± 0.04 | 0.42 ± 0.03 | 0.44 ± 0.02 |
|  | Median | 0.46 | 0.45 | 0.49 | 0.52 | 0.4 | 0.43 |
| **Direction Index<sup>2</sup>** | Mean +SE | 0.28 ± 0.03 | 0.3 ± 0.05 | 0.2 ± 0.02 | 0.35 ± 0.03 | 0.32 ± 0.03 | 0.29 ± 0.02 |
|  | Median | 0.22 | 0.2 | 0.2 | 0.33 | 0.33 | 0.28 |
| **Tuning Width (HWHH)<sup>3</sup>** | Mean +SE | 32.4 ± 2.2 | 35.4 ± 2.2 | 32.5 ± 1.3 | 35.7 ± 1.8 | 37.8 ± 2 | 35.5 ± 1.3 |
|  | Median | 24.6 | 34.3 | 31.5 | 35 | 34.3 | 34.3 |
| **F1/F0<sup>4</sup>** | Mean +SE | 0.33 ± 0.03 | 0.28 ± 0.04 | 0.33 ± 0.02 | 0.3 ± 0.03 | 0.23 ± 0.02 | 0.29 ± 0.02 |
|  | Median | 0.28 | 0.2 | 0.28 | 0.25 | 0.21 | 0.27 |

1. Normalized difference between the sum of responses to the preferred orientation vs. the sum of responses to the opposite orientation. 2. Normalized difference between the response to the preferred direction vs. that of the opposite direction. 3. Sharpness of direction tuning (deg) of responses (spikes/s) to drifting gratings moving in different directions. Values of sharpness of direction tuning correspond to the half-width at half-height of the maximum amplitude of tuning curve (HWHH). 4. F1: Amplitude of the first Fourier harmonic of V1 responses (spikes/s) to drifting grating stimulus; F0: Mean firing rate (spikes/s).

#### Sharpness of direction tuning

We evaluated the specificity of V1 neurons for direction of stimulus movement by measuring the sharpness of the direction tuning curve at half of the maximum amplitude of neuronal responses (firing rate). Direction tuning sharpness was not significantly influenced by rearing conditions at ∼P30 or ∼P60 (**Fig. 4C**, **Table 2**). V1 neurons across all ages and light conditions showed narrow direction tuning curves with a median half-width at half- height of the maximum response (HWHH) ranging from 20° to 40°. However, neurons from ∼P90 DR mice exhibited broader direction tuning than did neurons from ∼P90 NR mice (Mann-Whitney *U* test = 1724, *p* = 0.03). Thus, V1 neurons in dark reared mice compared to normally reared mice lost some of their specificity for stimulus movement direction as they reached adulthood.

#### Orientation selectivity

Stimulus orientation tuning in layer 2/3 V1 neurons was apparent in NR and DR mice by ∼P30, but the degree of selectivity was reduced at later ages by dark rearing. We assessed the orientation selectivity by calculating the orientation index (OI), which is the normalized difference between the sum of responses to the preferred orientation vs. the sum of responses to the opposite orientation. Across all V1 neurons analyzed for orientation selectivity, OI values did not show significant differences between rearing conditions at the different ages evaluated (∼P30: *t*-test = -1.2, *p* = 0.23; ∼P60: Mann-Whitney *U* test = 611, *p* = 0.4; ∼P90: Mann-Whitney *U* test = 1829, *p* = 0.15) (**Fig. 4D**, **Table 2**). However, a significant decrease in OI was observed between age groups of DR mice between ∼P30 and ∼P60 (Kruskal-Wallis 1-way ANOVA on ranks, H(2) = 8.9, *p* = 0.01, Dunn’s post hoc test, *p* < 0.05) and between ∼P30 and ∼P90: (Dunn’s post hoc test, *p* < 0.05), but not in NR mice:∼P30 vs. ∼P60 vs ∼P90: Kruskal-Wallis 1-way ANOVA on ranks, H(2) = 0.3, *p* = 0.9). In NR mice, from the total of V1 neurons used for orientation selectivity analysis, the percentage of single units with OI > 0.5 was 46% at ∼P30 (18/39 neurons), 42% at ∼P60 (12/28 neurons) and 48% at ∼P90 (32/66 neurons). In DR mice, the percentage of single units with OI > 0.5 was 53% at ∼P30 (24/45 neurons), 26% at ∼P60 (13/50 neurons) and 32% at ∼P90 (21/65 neurons). In sum, highly selective neurons were more frequently observed in adult NR than in adult DR mice. Overall, our results using gratings as stimuli show that in mouse V1 neurons there are tuning properties that require light exposure during postnatal development for optimal function including orientation selectivity and sharpness of direction tuning. However, others, including direction selectivity, arise independent of visual experience.

### The high proportion of V1 neurons showing non-linear responses was not affected by light rearing conditions

V1 neurons can be categorized as simple or complex by their responses, with simple (*aka* linear) neurons possessing spatially separate excitatory and inhibitory regions in their RFs and complex (*aka* non-linear) neurons exhibiting position invariance (Hubel & Wiesel, 1962). Here, we assessed the proportion of simple and complex neurons in V1 across ages and visual experience groups. Neurons in layer 2/3 that exhibited linear vs. non-linear responses were found to be present in similar proportions in NR and DR mice by Niell & Stryker (2008). In contrast, we found that 99% of the visually responsive neurons recorded exhibited nonlinear responses (F1/F0 < 1) across both age and rearing condition (F1/F0 median from 0.2 - 0.3) (**Fig. 4E**, **Table 2**). A difference in size and type of electrode used may account for the difference in proportions of single vs. complex neurons isolated across studies. V1 neurons did not exhibit significant changes in F1/F0 across age in either NR or DR mice (NR: Kruskal-Wallis 1-way ANOVA on ranks, H(2) = 5.4, *p* = 0.07; DR: Kruskal-Wallis 1-way ANOVA on ranks, H(2) = 5.7, *p* = 0.06), thus, the layer 2/3 V1 neurons included in this study were almost exclusively complex neurons in both groups, suggesting that DR had no effect on the proportions of simple and complex types of neuron.

### The suppressive effect of surround stimulation on V1 neurons was shaped by visual experience

Next, to evaluate the spatial arrangement of lateral inhibitory inputs of V1 layer 2/3 neurons and their effect on responses to light stimulation, we measured the evoked activity at the RF center (within which the spike rate of each single unit was highest) when different portions of the RF surround were stimulated. This was done using a combination of filled and ring-shaped gratings with a different external and internal radius (Gao et al., 2010; Popović et al., 2018). The effect of surround stimulation on the response magnitude of individual neurons was quantified as a stimulus size preference parameter (S*m*; see Methodology). *Sm* represents the degree and direction of modulation of the response at the RF center by surround stimulation. S*m* values are positive when stimulation of the surround monotonically increases the response at the RF center, including when the largest filled grating was used (surround facilitation), and negative when such stimulation reduces the response at the RF center (surround suppression), as defined by Popović et al. (2018). In mouse V1, we found bidirectional modulation of responses to different stimulus sizes across ages and rearing conditions (**Fig. 5A)**. Across all ages, V1 neurons exhibiting surround suppression were predominant in NR mice (∼P30 S*m* < 0: 89% (25/28 neurons); ∼P60 S*m* < 0: 100% (28/28 neurons); ∼P90 S*m* < 0: 95% (53/56 neurons)) (**Fig. 5B**, **Table 3**). In contrast, V1 in DR mice contained a lower percentage of neurons with surround suppression than did V1 in NR mice, across all age groups (∼P30: S*m* < 0: 81% (26/32 neurons), ∼P60 S*m* < 0: 90% (73/81 neurons) and ∼P90 S*m* < 0: 83% (96/115 neurons). DR mice at ∼P90 exhibited a significant increase in the proportion of neurons showing surround facilitation (S*m* > 0) in comparison to ∼P60 (10% at P60 and 17% at P90) (Kruskal-Wallis 1-way ANOVA on ranks, H(2) = 12.1, *p* = 0.002, Dunn’s post hoc test, *p* < 0.05), that was not seen in NR mice (Kruskal-Wallis 1-way ANOVA on ranks, H(2) = 2, *p* = 0.4). At ∼P90, V1 neurons in DR mice exhibited fewer negative and more positive *Sm* values than V1 neurons in NR mice (Mann-Whitney *U* test = 1961, *p* < 0.001). Thus, surround suppression can arise independent of visual experience, although the proportion of neurons showing suppression was reduced under dark rearing conditions, with DR mice exhibiting a higher proportion of neurons with stronger responses to stimuli of increasing size (surround facilitation neurons) than NR mice in adulthood.

**Figure 5.**
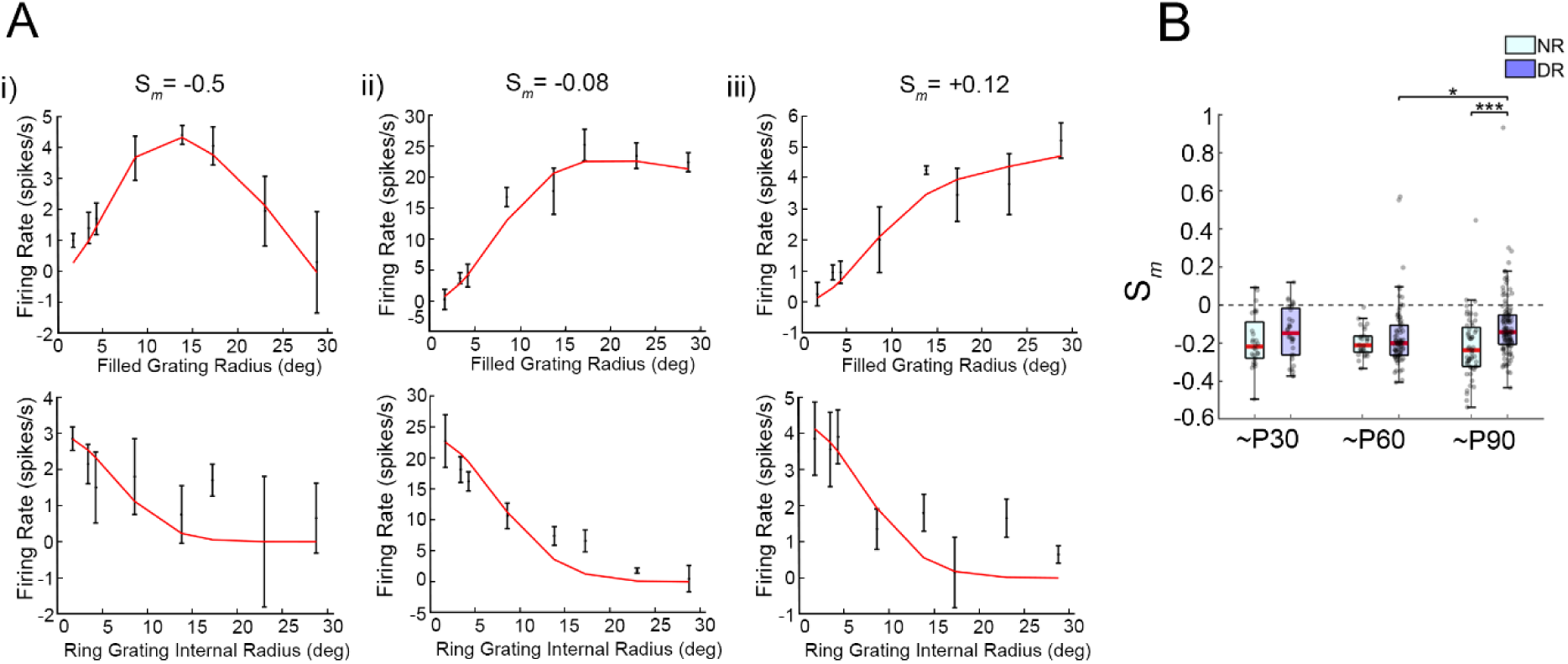
The development of the suppressive effect of surround stimulation on V1 neurons is dependent on visual experience. **(A)** Examples of single unit tuning curves showing mean evoked firing rate ± S.E.M to 8 filled gratings **(upper)** and 8 ring-shaped gratings **(lower)** of different sizes. From left to right, neurons showing suppression of their responses to stimulation of the center of the RF (size modulation (S*_m_*) <0, left), neurons showing no modulation of their responses (S*_m_* ∼0, middle) and neurons showing surround facilitation (S*_m_*>0, right) are shown. **(B and C)** Each box shows the median (red lines) and 25th and 75th percentiles. Whiskers correspond to minimum and maximum values within a 1.5x interquartile range. **(B)** Quantification of size modulation (S*_m_*) values across ages and light conditions. **(C)** RF radius of single unit responses to filled and ring-shaped gratings across ages and rearing conditions *p<0.05, ***p<0.001.

**Table 3.**
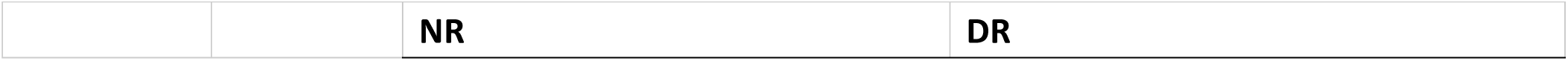
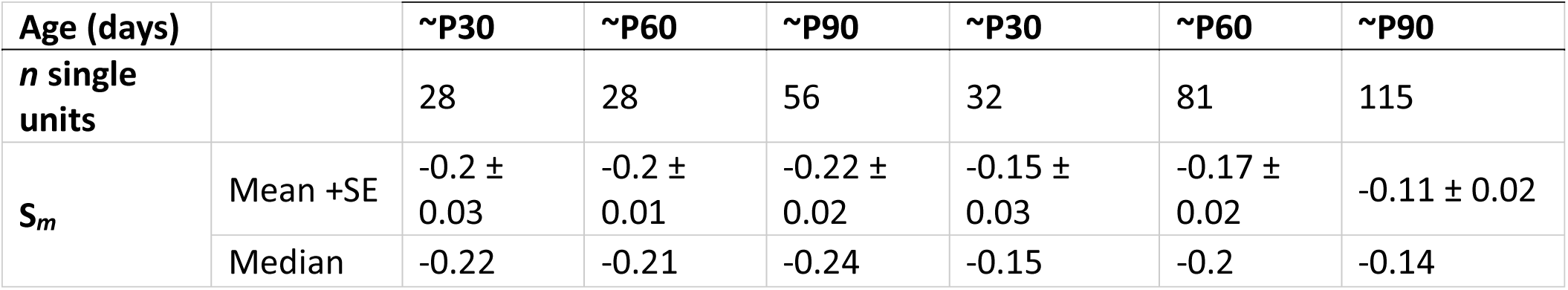
Stimulus size tuning properties assayed with filled and ring-shape drifting gratings.

| Age (days) |  | ~P30 | ~P60 | ~P90 | ~P30 | ~P60 | ~P90 |
| --- | --- | --- | --- | --- | --- | --- | --- |
| ***n* single units** |  | 28 | 28 | 56 | 32 | 81 | 115 |
| ***S<sub>m</sub>*** | Mean +SE | -0.2 ± 0.03 | -0.2 ± 0.01 | -0.22 ± 0.02 | -0.15 ± 0.03 | -0.17 ± 0.02 | -0.11 ± 0.02 |
|  | Median | -0.22 | -0.21 | -0.24 | -0.15 | -0.2 | -0.14 |

### Visual experience caused V1 neurons to develop clear RF boundaries with stronger surround suppression compared to mice reared in prolonged darkness

Our group has shown that in Syrian hamsters the maintenance of refined RFs in SC of NR but not DR hamsters is largely due to the maintenance of a strong inhibitory surround (Carrasco et al., 2005; Carrasco et al., 2011). To investigate the connection between RF size and surround suppression, we calculated the suppression ratio (SR, see Methodology) of individual V1 neurons in mice. Distinct from *Sm*, SR evaluates the modulation of the RF responses by center and surround stimulation using filled, drifting gratings of increasing size (**Fig. 6A**) (Popović et al., 2018). Neurons showing neuronal responses that are weakly suppressed by surround stimulation, as well as those in which the neuronal response was not increased or was facilitated, exhibit an SR close or equal to 1. In contrast, neurons exhibiting strong surround suppression reduced response levels as the stimulus enlarged (SR <1). To ensure greater accuracy of the results, the SR was calculated only for the V1 neurons that showed clear, elliptical RFs when a flashing white square was used (**Fig. 6B**). At P30, no differences in SR were found between NR and DR groups (∼P30 NR vs. DR: Mann-Whitney *U* test = 621, *p* =0.9). In neurons from NR mice, suppression by surround stimulation increased with age (SR decreased): between ∼P30 vs. ∼P60 (Kruskal-Wallis 1-way ANOVA on ranks H(2) = 22.3, *p* < 0.001, Dunn’s post hoc test, *p* < 0.05) and between ∼P30 vs. ∼P90 (Dunn’s post hoc test, *p* < 0.05) (**Fig. 6C**, **Table 4**). In contrast, neurons in DR mice exhibited a weaker suppressive surround effect (SR values close to 1) than neurons in NR mice, with no change across ages: ∼P30 vs. ∼P60 vs. ∼P90 (Kruskal-Wallis One-way ANOVA on ranks, H(2) = 1.96, *p* = 0.4). At older ages, SR values were significantly smaller in NR mice compared to DR mice (∼P60 NR vs DR: Mann-Whitney *U* test = 772, *p* = 0.003; ∼P90 NR vs. DR: Mann-Whitney *U* test = 198, *p* < 0.001), due to greater surround suppression in the NR mice. Consistent with this we found that responses of V1 neurons in NR animals to drifting gratings of small size increased with age (**Fig. 6D**, **Table 4**). At P30, no differences in stimulus size preference were found between NR and DR groups (∼P30 NR vs. ∼P30 DR: Mann-Whitney *U* test = 541, *p* = 0.3). However, NR animals at both ∼P60 and ∼P90 showed stronger responses to smaller stimuli than they did at ∼P30: between ∼P30 vs. ∼P60 (Kruskal-Wallis 1-way ANOVA on ranks, H(2) = 18.8, *p* <0.001, Dunn’s post hoc test, *p* < 0.05) and between ∼P30 vs. ∼P90 (Dunn’s post hoc test, *p* < 0.05). A significant preference for smaller stimuli in NR in comparison to DR animals was observed at ∼P60 (NR vs. DR: Mann-Whitney *U* test = 843, *p* = 0.01) and ∼P90 (NR vs. DR: Mann-Whitney *U* test = 541, *p* < 0.001). In contrast to changes in V1 neuron stimulus size preference in NR mice, no significant differences in the preferred stimulus size were found across ages in DR mice (∼P30 vs. ∼P60 vs. ∼P90: Kruskal-Wallis 1-way ANOVA on ranks, H(2) = 2, *p* = 0.4). Thus, V1 neurons in mice reared in normal light conditions develop stronger surround suppression than neurons in those raised in prolonged darkness, resulting in a preference for a smaller stimulus size compared to DR mice.

**Figure 6.**
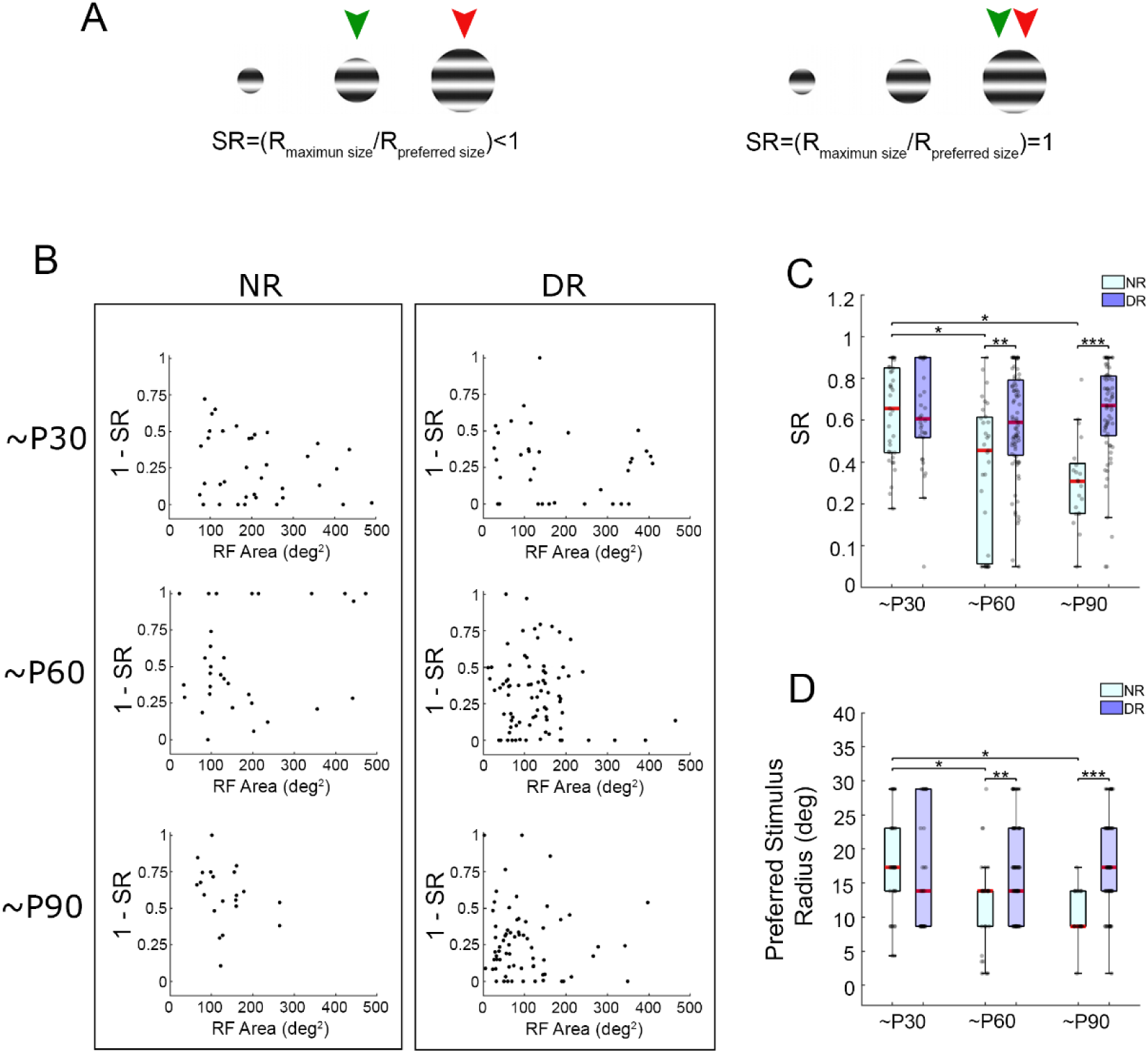
Surround suppression in V1 neurons became stronger with age only in visually experienced mice. **(A)** Suppression ratio (SR). V1 neuron responses to the grating size giving the maximum response **(green arrowhead)** and the largest grating used **(red arrowhead)** were used to calculate SR. (**B**) Reciprocal SR values (1-SR) calculated from V1 neurons showing defined RF areas for RF size assay, plotted against RF area (deg^2^) of NR and DR mice at different ages. Dashed line at reciprocal SR value of 0.5 is shown for each plot. (**C**) Boxplot of SR values of NR and DR mice at different ages. **D**) Boxplot showing the radius of the filled grating stimulus triggering the maximum response (preferred stimulus radius) of NR and DR mice at different ages. Each box shows the median **(red lines)** and 25th and 75th percentiles. Whiskers correspond to minimum and maximum values within a 1.5x interquartile range. *p<0.05, **p<0.01, ***p<0.001.

**Table 4.** Stimulus size tuning properties of V1 neurons assayed with a flashing white square showing clear RF boundaries.

|  |  | NR |  |  | DR |  |  |
| --- | --- | --- | --- | --- | --- | --- | --- |
| Age (days) |  | ~P30 | ~P60 | ~P90 | ~P30 | ~P60 | ~P90 |
| ***n* single units** |  | 37 | 31 | 22 | 34 | 79 | 67 |
| **Suppression Ratio (SR)** | Mean +SE | 0.74 ± 0.04 | 0.46 ± 0.06 | 0.41 ± 0.04 | 0.72 ± 0.04 | 0.68 ± 0.03 | 0.74 ± 0.77 |
|  | Median | 0.76 | 0.56 | 0.41 | 0.71 | 0.69 | 0.03 |
| **Preferred Stimulus Radius (deg)** | Mean +SE | 17.5 ± 1.2 | 12.4 ± 1.1 | 10.6 ± 0.7 | 16.2 ± 1.5 | 16.6 ± 0.8 | 17.5 ± 0.8 |
|  | Median | 17.3 | 13.8 | 8.6 | 13.8 | 15.6 | 17.3 |

### Visual experience caused V1 neurons to develop stronger responses to stimulus size matching their RF sizes compared to mice reared in prolonged darkness

Most neurons in the primary visual cortex of primates, carnivores, and rodents respond optimally to stimuli matching their RF size. Stimuli larger than the RF can decrease the response to stimuli at the RF center due to surround inhibition (Allman et al., 1985; Cavanaugh et al., 2002; Adesnik et al., 2012; Angelucci et al., 2017; Samonds et al., 2017). Surround facilitation can also occur, in which the maximum response is evoked by stimuli larger than the RF center (Gao et al. 2010; Popović et al., 2018 and this study). We examined the relationship between center and surround RF areas using a flashing white square (see **Fig. 1B**) and determined the size of filled grating stimulus that triggered the maximum response in V1 neurons of NR and DR mice (**Fig. 7A**). In NR mice, the percentage of V1 neurons exhibiting a facilitatory response to stimuli larger in size than the RF decreased with age, and was 81% at ∼P30, 80% at ∼P60 and 57% at ∼P90. In contrast, in DR mice, the percentage of neurons exhibiting this trait was 78% at ∼P30, 96% at ∼P60 and 95% at ∼P90 (**Table 5**). Thus, the proportion of neurons exhibiting maximum responses to stimuli larger than their RF size decreased with age under normal rearing conditions but increased with age in DR mice.

**Figure 7.**
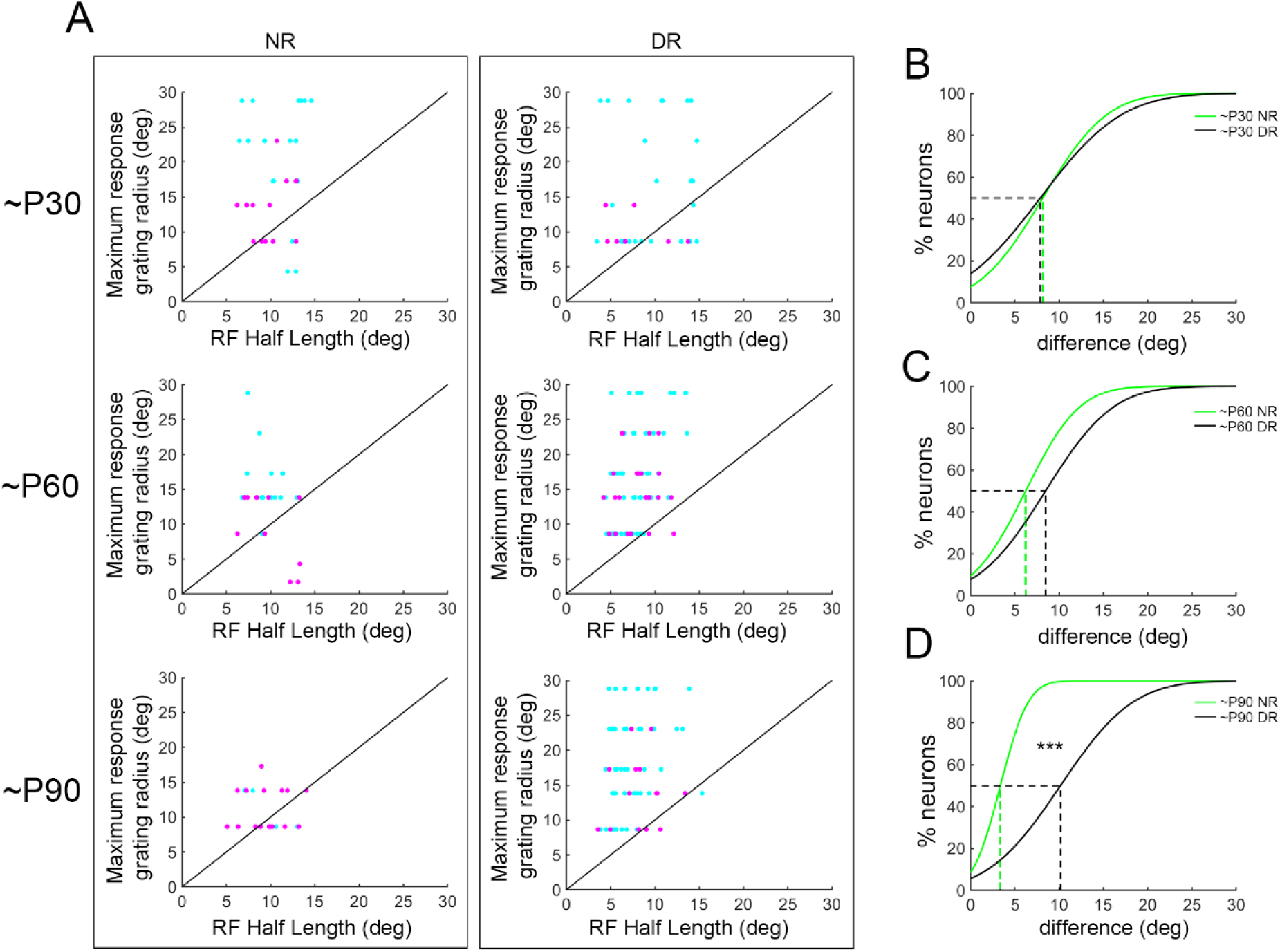
V1 neurons in NR mice developed optimal neuronal responses to stimuli matching their RF size, but a lack of visual experience resulted in a preference for stimuli larger than the RF. **A)** For each V1 neuron showing a measured RF using flashing white squares, the RF half-length (degrees) was compared against the filled grating radius (i.e. stimulus size) triggering the maximum response (maximum response grating radius (degrees)). The proportion of neurons responding optimally to stimuli larger than their RF decreases with age in normal conditions, but increases in mice raised in darkness. Black line corresponds to the identity line (slope *m*=1). For visualization, neurons showing a strong surround inhibition (SR<0.6, magenta) and weak surround inhibition (SR>0.6, cyan) are shown. **(B-D)** Cumulative probability showing the distribution of the differences (deg) between the RF half-length and the filled grating radius that triggers a maximum response for V1 neurons of NR and DR mice (see Methods section). Distributions were calculated at∼P30 (B), ∼P60 **(C)** and ∼P90 **(D)**. Dashed lines indicate the median values for each distribution. V1 neurons developed optimal responses to stimuli that equal the RF size in NR mice, but in DR mice neurons responded optimally to stimuli that were larger than the RF. This difference between NR and DR was highly significant at P90. ***p<0.001.

**Table 5.** Percentages of V1 neurons exhibiting modulation of their responses by stimulus size.

|  | NR |  |  | DR |  |  |
| --- | --- | --- | --- | --- | --- | --- |
| Age (days) | ~P30 | ~P60 | ~P90 | ~P30 | ~P60 | ~P90 |
| ***n* single units** | 36 | 25 | 21 | 33 | 78 | 65 |
| % of single units with optimal stimulus size* > RF size | 81 | 80 | 57 | 79 | 96 | 95 |
| % of single units with optimal stimulus size* < RF size | 19 | 20 | 42 | 21 | 4 | 5 |
| % of single units with SR** < 0.6 | 36 | 48 | 81 | 24 | 33 | 21 |

\*Stimulus size (filled grating radius) triggering maximum response
\*\* SR = Suppression ratio

**Figure 8.**
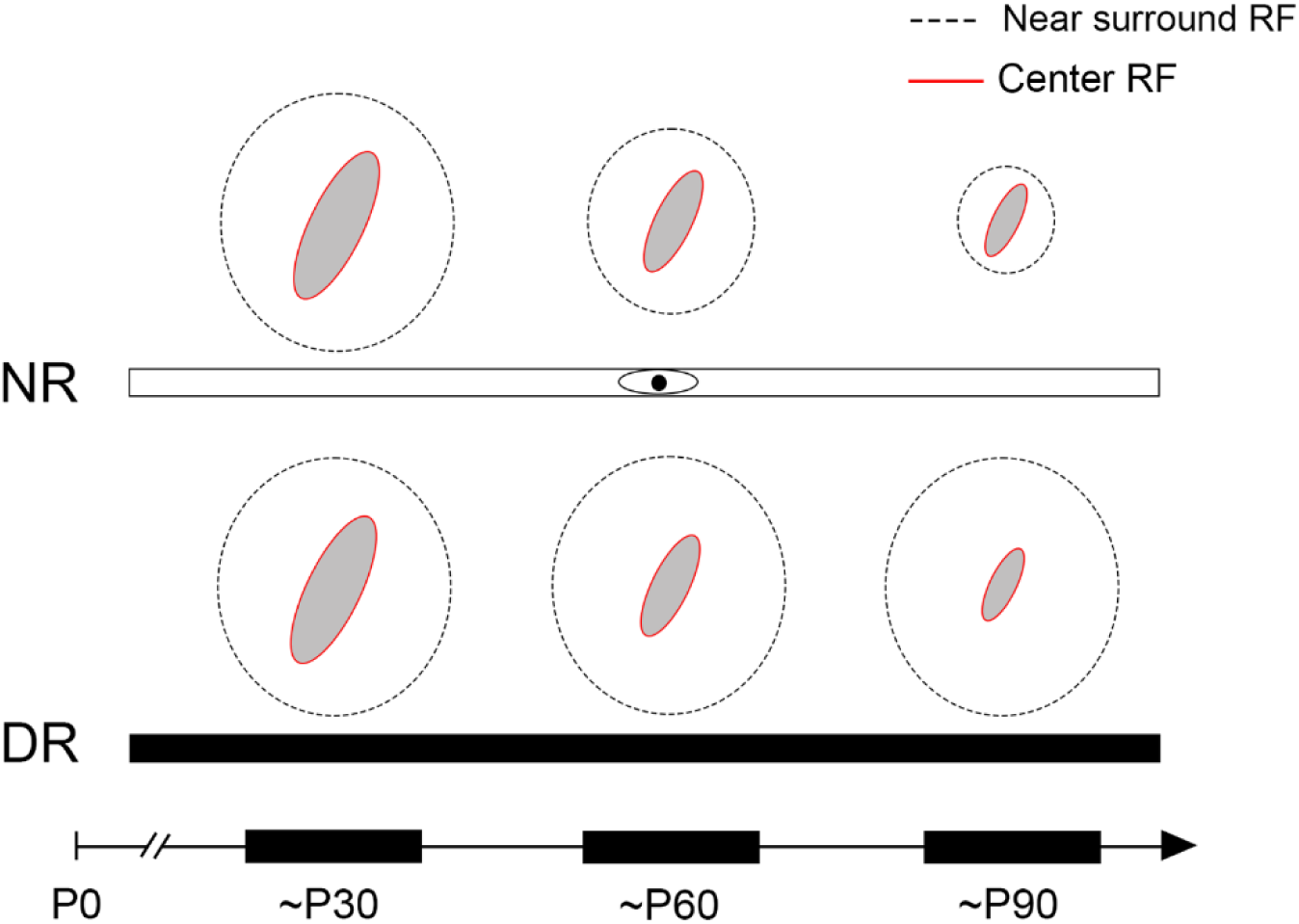
Refinement of RF center size in V1 layer 2/3 neurons across postnatal development was similar between mice reared in normal light conditions and those reared in continuous darkness. At early ages of both NR and DR mice V1 neurons presented with flashing white squares that were moved randomly across the screen exhibited large RF centers. Across later ages the majority were refined to a smaller size. In contrast, the near surround of RFs, defined by the size of the filled grating stimulus that triggered the maximum response of V1 neurons, was refined in 57% NR mice at ∼P90 but did not change in size across ages in 95% of neurons in DR mice at the same age.

Next, we evaluated how the size of the stimulus evoking the maximum response varied with respect to the RF size of V1 neurons by assessing the distribution of the differences between classical RF size and the stimulus size at which neurons responded maximally across ages in NR and DR mice (**Fig. 7B-D**; see Methodology). At ∼P30, the distribution of the RF size and the stimulus size causing the maximum response (RF size + excitatory surround extent) between NR and DR mice did not reach a significant difference (median ∼P30 NR = 8.1 deg, median ∼P30 DR = 7.8 deg; Kolmogorov-Smirnov test, *p* = 0.1) (**Fig. 7B**). There was also a nearly significant difference at ∼P60 (median ∼P60 NR = 6.2 deg, median ∼P60 DR = 8.4 deg; Kolmogorov-Smirnov test, *p* = 0.07) (**Fig. 7C**). However, at ∼P90, the difference between the RF size and the size of the stimulus triggering the maximum response was significantly larger in V1 neurons of DR mice than in V1 neurons of NR mice (median ∼P90 NR = 3.4 deg, median ∼P90 DR = 10.1 deg; Kolmogorov-Smirnov test, *p* = 9.2 x10^-6^) (**Fig. 7D**). These results show that with age, V1 neurons in NR mice develop optimal responses to stimuli with a diameter that better matches their RF size, but the development of this trait is negatively affected by dark rearing.

In summary, although a suppressive influence of the surround on V1 neuron responses existed in both normal and dark rearing conditions, the degree of suppression increased with age only in visually experienced mice. This was accompanied by stronger responses to stimuli better matching their RF sizes, compared to mice reared in prolonged darkness. These results support the interpretation that dark rearing causes a reduction in lateral inhibition and/or an increase of near surround facilitation, despite having no effect on the classical RF size or the presence of orientation selectivity. Future studies focusing on the molecular mechanisms involved in the excitatory and inhibitory regulation of V1 neuron responses in mice and the role of visual experience in their establishment, similar to previous studies in hamsters (Carrasco et al., 2011; Mudd et al., 2019), would clarify this point.

## DISCUSSION

The results from the present study on the role of visual experience in the development and maintenance of several receptive field properties in primary visual cortex of lab mice provide support for the hypothesis that diel activity pattern rather than phylogeny is the determining factor. Contrary to what has been reported previously in some other mammalian species, we found that RFs of layer 2/3 V1 neurons in lab mice reared in complete darkness attain and maintain refinement of their excitatory center. In addition, as reported previously (Rochefort et al., 2011), we observed that visual experience was not necessary for development of stimulus direction selectivity in mice. However, other response properties, including orientation selectivity and surround suppression, required light exposure to develop normally. Of course, if visible objects are available, mice can use them to influence behavior (e.g., Zhang et al., 2026). Compared with our previous findings in hamsters (Carrasco et al., 2005; Carrasco and Pallas, 2006; Balmer and Pallas, 2015) and those from others in additional mammalian species (see Pallas, 2017, for review) our current results argue that the environment and ecological niche of each species over evolutionary time is more influential than phylogeny on the necessity for visual experience during development and adulthood.

We quantified the area and half-length of RFs in V1 of mice throughout postnatal development and into adult ages in both NR and DR groups. As expected from studies on juveniles of other mammalian species including rats (Fagiolini et al., 1994), hamsters (Carrasco et al., 2005; Balmer and Pallas, 2015), ferrets (Popović et al., 2018; Fernandez-Aburto et al., 2024), cats (Blakemore and Van Sluyters, 1975; Hirsch, 1985), and monkeys (Jacobs and Blakemore, 1988; Kiorpes, 2015), we found that RFs of layer 2/3 neurons in V1 of NR juvenile mice (∼P30) were large in comparison to RFs in adults. The same was true for juvenile DR mice. RFs were significantly larger at ∼P45 than in adults, both in normal mice and after short term DR (Gianfranceschi et al., 2003; Ko et al., 2013; 2014), but DR mice have larger RFs than NR mice (Gianfranceschi et al., 2003). At ∼P60 (sexual maturity) we found that there was still no significant difference in RF size between NR and DR mice. We reported similar findings previously in both SC and V1 of hamsters; dark rearing until ∼P60 had no effect. However, continued dark rearing in hamsters past ∼P60 led to a loss of RF refinement and a degradation in visual perception in both SC and V1 (Carrasco et al., 2005; Balmer and Pallas, 2015). In DR mice, we found that classical, excitatory RFs are refined to a normal extent by ∼P90, arguing that the developmental refinement and adult maintenance of RF size in V1 layer 2/3 neurons occurs without any visual experience. Given that hamsters are crepuscular in the wild, but mice are nocturnal, this result supports the hypothesis that ecological niche and diel activity pattern are more influential than phylogeny for the process of receptive field refinement and maintenance.

Spontaneous activity plays a crucial role in the early development of visual cortical response properties in rats (Benevento et al., 1992), ferrets (Chapman and Stryker, 1993; Chiu and Weliky, 2001; White et al., 2001; Li et al., 2006; Li et al., 2008; Griswold and Van Hooser, 2025) and cats (Huttenlocher, 1967; Callaway and Katz, 1990; Arieli et al., 1995; Bringuier et al., 1997; Tanaka et al., 2004; Murphy and Miller, 2009; Ch’ng and Reid, 2010). From our results, we conclude that spontaneous activity in DR mice is sufficient to drive RF refinement of V1 neurons under continuous dark rearing. We reported a similar phenomenon in dark reared hamsters; spontaneous activity alone can drive the refinement (although not the maintenance) of their RFs in both V1 and superior colliculus (Carrasco et al., 2005; Balmer and Pallas, 2015). Thus, in the absence of light exposure, spontaneous activity is sufficient for the complete refinement of RF size in rodents exhibiting non-diurnal visual habits. Evoked responses to visual stimulation are necessary only to maintain RF refinement in both V1 (Balmer and Pallas, 2015) and SC (Carrasco et al., 2005) neurons in adult hamsters.

In this study, evoked spike rates of V1 neurons to light stimuli differed across ages in mice, being stronger in NR V1 neurons than in DR V1 neurons at ∼P90. We suggest that in mice, the lower evoked response levels observed in DR compared to NR adults might be related to the reduced sensitivity of neurons in adulthood resulting from a general decline in response levels after long periods of dark rearing.

In adult mice, most of the neurons in V1 layer 2/3 are sharply tuned to stimulus orientation, and a lower proportion are tuned to stimulus movement direction (Drager, 1975; Mangini and Pearlman, 1980; Niell and Stryker, 2008). We found a similar trend in adults, with nearly half of the neurons in ∼P90 NR mice exhibiting OI > 0.5, consistent with what was found by Gao and colleagues (2010). V1 neurons in our dark reared mice did not change their degree of selectivity for direction. In contrast, direction tuning was broader in dark-reared adults. Selectivity for orientation and the proportion of highly selective neurons for orientation (OI>0.5) decreased with age in DR cases (Kuhlman et al., 2011; Rochefort et al., 2011). Our results differ from findings in ferrets (Popović et al., 2018), cats (Leventhal and Hirsch, 1980) and rats (Fagiolini et al., 1994), in which direction selectivity of V1 neurons requires visual experience (Li et al., 2006; 2008; Elstrott and Feller, 2009). Therefore, visual experience in mice drives the development and maintenance of selectivity for stimulus orientation and the sharpening of direction tuning in V1 neurons across the ages evaluated, but not the preferred directions across the population of neurons recorded.

Unexpectedly, we did not find simple V1 neurons in mice across ages and rearing conditions. In contrast, Niell & Stryker (2008) reported in adult mice that V1 neurons exhibiting linear and nonlinear responses were present in equivalent proportions. One possible explanation for the presence of only complex cells found in our study is that, in contrast to Niell & Stryker (2008), our analysis of linearity considered V1 responses only at the preferred direction of grating stimulus, but not at the preferred spatial frequency. Thus, most of the V1 neurons found in our study exhibit properties typical of complex-type V1 neurons.

In this study, we observed that the suppressive effect of surround stimulation increases with age in V1 of mice raised in normal but not in dark conditions. In V1, visual experience is necessary for developmental strengthening of inhibition (Morales et al., 2002; Katagiri et al., 2007). In hamster SC (Rhoades and Chalupa, 1978; Carrasco et al., 2005; Mudd et al., 2019) and visual cortex (Balmer and Pallas, 2015) the presence of a strong inhibitory surround restricts RF size. Prolonged DR into adulthood in hamsters reduces the strength of the inhibitory surround in SC, which would result in a failure to maintain refined RFs (Carrasco et al., 2011; Mudd et al., 2019). In this study, although a suppressive effect of surround stimulation over V1 neuron responses was seen at early ages, the effect became stronger with age in NR, but not in DR subjects.

We found that a high percentage of V1 neurons in juvenile NR mice exhibited facilitatory response modulation in reaction to surround stimulation. By adulthood, however, RFs are restricted in response area by strong surround suppression, with preferred stimulus sizes better matching their RF size. RF center is defined as the excitatory central portion and the surround as the region outside of it. Both center and surround areas were visualized depending on the stimulus used, with sparse noise (flashing white squares) showing the classical RF, and stationary gratings of different spatial frequencies showing multiple and elongated on and off subregions of RFs (Yeh et al., 2009). It illustrates that RF shape is not stimulus-invariant and suggests that intracortical interactions in layer 2/3 neurons shape RF properties. Interestingly, Angelucci and Bresloff (2006) observed a facilitatory near surround between the excitatory RF center of V1 neurons and the far surround (usually inhibitory but can also be excitatory) in monkeys. The use of a flashing square stimulus allowed the calculation of the excitatory RF center with no influences of its modulatory surround (Angelucci and Bressloff, 2006; Fang et al., 2021). Filled gratings of different spatial frequency instead allow the calculation of the excitatory RF center and its modulatory surround, including the facilitatory near surround (Angelucci and Bressloff, 2006; Gao et al., 2010; Popović et al., 2018, and the present study). We thus suggest that in mice, V1 development under normal conditions involves the reduction of the facilitatory near surround of V1 RFs through the increase of surround inhibition with age, but its maintenance under prolonged darkness conditions. Excitatory RF centers, in contrast, were reduced in size independent of visual experience.

In conclusion, we have shown that the refinement and maintenance of RF size and the selectivity for direction of movement in V1 layer 2/3 neurons occur independent of visual experience in mice. In contrast, orientation and stimulus size tuning require visual experience. Our results support our hypothesis that whether visual system development depends on visual experience is determined by ecological niche with respect to diel activity pattern and nest location and thus access to light stimulation rather than by phylogeny (see also Campi and Krubitzer, 2010). Laboratory rodents exhibiting crepuscular or nocturnal sleep/wake habits and developing in underground nests rely more heavily on olfactory and somatosensory cues than on their limited visual perception for survival. Therefore, light exposure before weaning is less crucial for them to construct visual circuits than it is for diurnal species with well-developed visual systems. Finally, our results reveal the limits of the mouse as a model for human vision research, prompting the need for more suitable animal models, such as diurnal rodents, for future research on visual disorders.

## Acknowledgments

We would like to thank members of the Pallas lab for technical support, and the animal care staff at UMass-Amherst for their dedicated attention to the mice in our facility. Prof. Gonzalo Marín and Dr. Natalia Márquez provided valuable help with their comments on the manuscript. Thanks also go to Prof. Stephen van Hooser for providing some of the functions used for data analysis. Support for this work was provided by UMass-Amherst startup funds and National Science Foundation funding (IOS-1656838, IOS-2029980) to S.L.P.

## REFERENCES

Adesnik H, Bruns W, Taniguchi H, Huang ZJ, Scanziani M (2012) A neural circuit for spatial summation in visual cortex. Nature 490:226–231.

Allman J, Miezin F, McGuinness E (1985) Stimulus specific responses from beyond the classical receptive field: neurophysiological mechanisms for local-global comparisons in visual neurons. Annu Rev Neurosci 8:407–430.

Angelucci A, Bressloff PC (2006) Contribution of feedforward, lateral and feedback connections to the classical receptive field center and extra-classical receptive field surround of primate V1 neurons. Prog Brain Res 154(A):93–120

Angelucci A, Bijanzadeh M, Nurminen L, Federer F, Merlin S, Bressloff PC (2017) Circuits and mechanisms for surround modulation in visual cortex. Ann Rev Neurosci 40:425–451.

Arieli A, Shoham D, Hildesheim R, Grinvald A (1995) Coherent spatiotemporal patterns of ongoing activity revealed by real-time optical imaging coupled with single-unit recording in the cat visual cortex. J Neurophysiol 73:2072–2093.

Balmer TS, Pallas SL (2015) Refinement but not maintenance of visual receptive fields is independent of visual experience. Cereb Cortex 25:904–917.

Berardi N, Pizzorusso T, Maffei L (2000) Critical periods during sensory development. Curr Opin Neurobiol 10:138.

Berman D, Daw NW (1977) Comparison of the critical periods for monocular and directional deprivation in cats. J Physiol (Lond) 265:249–259.

Blakemore C, Van Sluyters RC (1975) Innate and environmental factors in the development of the kitten’s visual cortex. J Physiol 248:663–716.

Blakemore C, Garey LJ, Vital-Durand F (1978) The physiological effects of monocular deprivation and their reversal in the monkey’s visual cortex. J Physiol 283:223–262.

Bringuier V, Frégnac Y, Baranyi A, Debanne D, Shulz DE (1997) Synaptic origin and stimulus dependency of neuronal oscillatory activity in the primary visual cortex of the cat. J Physiol 500(3):751–774.

Callaway EM, Katz LC (1990) Emergence and refinement of clustered horizontal connections in cat striate cortex. J Neurosci 10:1134–1153.

Campi KL, Krubitzer L (2010) Comparative studies of diurnal and nocturnal rodents: Differences in lifestyle result in alterations in cortical field size and number. J Comp Neurol 518:4491–4512.

Carandini M, Ferster D (2000) Membrane potential and firing rate in cat primary visual cortex. J Neurosci 20:470–484.

Carrasco MM, Pallas SL (2006) Early visual experience prevents but cannot reverse deprivation-induced loss of refinement in adult superior colliculus. Vis Neurosci 23:845–852.

Carrasco MM, Razak KA, Pallas SL (2005) Visual experience is necessary for maintenance but not development of receptive fields in superior colliculus. J Neurophysiol 94:1962–1970.

Carrasco MM, Mao YT, Balmer TS, Pallas SL (2011) Inhibitory plasticity underlies visual deprivation-induced loss of receptive field refinement in the adult superior colliculus. Eur J Neurosci 33:58–68.

Cavanaugh JR, Bair W, Movshon JA (2002) Nature and interaction of signals from the receptive field center and surround in macaque V1 neurons. J Neurophysiol 88:2530–2546.

Ch’ng YH, Reid RC (2010) Cellular imaging of visual cortex reveals the spatial and functional organization of spontaneous activity. Front Integr Neurosci 4.

Chapman B, Stryker MP (1993) Development of orientation selectivity in ferret visual cortex and effects of deprivation. J Neurosci 13:5251–5262.

Chapman B, Stryker MP, Bonhoeffer T (1996) Development of orientation preference maps in ferret primary visual cortex. JNeurosci 16:6443–6453.

Chapman B, Godecke I, Bonhoeffer T (1999) Development of orientation preference in the mammalian visual cortex. J Neurobiol 41:18–24.

Chen X-j, Rasch MJ, Chen G, Ye C-q, Wu S, Zhang X-h (2014) Binocular input coincidence mediates critical period plasticity in the mouse primary visual cortex. J Neurosci 34:2940–2955.

Chiu C, Weliky M (2001) Spontaneous activity in developing ferret visual cortex in vivo. J Neurosci 21:8906–8914.

Drager UC (1975) Receptive fields of single cells and topography in mouse visual cortex. J Comp Neurol 160:269–290.

Dräger UC, Olsen JF (1980) Origins of crossed and uncrossed retinal projections in pigmented and albino mice. J Comp Neurol 191:383–412.

Elstrott J, Feller MB (2009) Vision and the establishment of direction-selectivity: a tale of two circuits. Curr Opin Neurobiol 19:293–297.

Espinosa JS, Stryker Michael P (2012) Development and plasticity of the primary visual cortex. Neuron 75:230–249.

Fagiolini M, Pizzorusso T, Berardi N, Domenici L, Maffei L (1994) Functional postnatal development of the rat primary visual cortex and the role of visual experience: Dark rearing and monocular deprivation. Vision Res 34:709–720.

Fang Q, Li Y-t, Peng B, Li Z, Zhang LI, Tao HW (2021) Balanced enhancements of synaptic excitation and inhibition underlie developmental maturation of receptive fields in the mouse visual cortex. J Neurosci 41:10065–10079.

Fernandez-Aburto P, Sudana K, Beech E, Suarez Casanova V, Griswold SV, van Hooser SD, Pallas SL (2024) Maintenance but not refinement of receptive field size in ferret primary visual cortex requires visual experience. In: Society for Neuroscience Annual Meeting PSTR380.311. Chicago, IL: Society for Neuroscience.

Gandhi SP, Yanagawa Y, Stryker MP (2008) Delayed plasticity of inhibitory neurons in developing visual cortex. Proc Natl Acad Sci USA 105:16797–16802.

Gao E, DeAngelis GC, Burkhalter A (2010) Parallel input channels to mouse primary visual cortex. J Neurosci 30:5912–5926.

Gianfranceschi L, Siciliano R, Walls J, Morales B, Kirkwood A, Huang ZJ, Tonegawa S, Maffei L (2003) Visual cortex is rescued from the effects of dark rearing by overexpression of BDNF. Proc Natl Acad Sci USA 100:12486–12491.

Griswold SV, Van Hooser SD (2025) Premature vision drives aberrant development of response properties in primary visual cortex. Elife 14: RP106513.

Hirsch HVB (1985) The role of visual experience in the development of cat striate cortex. Cell Molec Neurobiol 5:103–121.

Hofer SB, Mrsic-Flogel TD, Bonhoeffer T, Hübener M (2006) Lifelong learning: ocular dominance plasticity in mouse visual cortex. Curr Opin Neurobiol 16:451–459.

Huang ZJ, Kirkwood A, Pizzorusso T, Porciatti V, Morales B, Bear MF, Maffei L, Tonegawa S (1999) BDNF regulates the maturation of inhibition and the critical period of plasticity in mouse visual cortex. Cell 98:739–755.

Hubel DH, Wiesel TN (1962) Receptive fields, binocular interaction and functional architecture in the cat’s visual cortex. J Physiol (Lond) 160:106–154.

Hubel DH, Wiesel TN (1970) The period of susceptibility of the physiological effects of unilateral eye closure in kittens. J Physiol (London) 206:419–436.

Hübener M (2003) Mouse visual cortex. Curr Opin Neurobiol 13:413–420.

Huttenlocher PR (1967) Development of cortical neuronal activity in the neonatal cat. Exptl Neurol 17:247–262.

Jacobs DS, Blakemore C (1988) Factors limiting the postnatal development of visual acuity in the monkey. Vision Res 28:947–958.

Kang E, Durand S, LeBlanc JJ, Hensch TK, Chen C, Fagiolini M (2013) Visual acuity development and plasticity in the absence of sensory experience. J Neurosci 33:17789–17796.

Katagiri H, Fagiolini M, Hensch TK (2007) Optimization of somatic inhibition at critical period onset in mouse visual cortex. Neuron 53:805–812.

Kiorpes L (2015) Visual development in primates: Neural mechanisms and critical periods. Dev Neurobiol 75:1080–1090.

Ko H, Mrsic-Flogel TD, Hofer SB (2014) Emergence of feature-specific connectivity in cortical microcircuits in the absence of visual experience. J Neurosci 34:9812–9816.

Ko H, Cossell L, Baragli C, Antolik J, Clopath C, Hofer SB, Mrsic-Flogel TD (2013) The emergence of functional microcircuits in visual cortex. Nature 496:96–100.

Kuhlman SJ, Tring E, Trachtenberg JT (2011) Fast-spiking interneurons have an initial orientation bias that is lost with vision. Nature Neurosci 14:1121–1123.

LeVay S, Wiesel TN, Hubel DH (1980) The development of ocular dominance columns in normal and visually deprived monkeys. J Comp Neurol 191:1.

Leventhal AG, Hirsch HV (1980) Receptive-field properties of different classes of neurons in visual cortex of normal and dark-reared cats. J Neurophysiol 43:1111–1132.

Li Y, Fitzpatrick D, White LE (2006) The development of direction selectivity in ferret visual cortex requires early visual experience. Nature Neurosci 9:676–681.

Li Y, Van Hooser SD, Mazurek M, White LE, Fitzpatrick D (2008) Experience with moving visual stimuli drives the early development of cortical direction selectivity. Nature 456:952–956.

Mangini NJ, Pearlman AL (1980) Laminar distribution of receptive field properties in the primary visual cortex of the mouse. J Comp Neurol 193:203–222.

Mazurek M, Kager M, Van Hooser SD (2014) Robust quantification of orientation selectivity and direction selectivity. Frontiers in neural circuits 8:92.

Morales B, Choi S-Y, Kirkwood A (2002) Dark rearing alters the development of GABAergic transmission in visual cortex. J Neurosci 22:8084–8090.

Mower GD (1991) The effect of dark rearing on the time course of the critical period in cat visual cortex. Devel Brain Res 58:151–158.

Mudd DB, Balmer TS, Kim SY, Machhour N, Pallas SL (2019) TrkB activation during a critical period mimics the protective effects of early visual experience on perception and the stability of receptive fields in adult superior colliculus. J Neurosci 39:4475–4488.

Murphy BK, Miller KD (2009) Balanced amplification: a new mechanism of selective amplification of neural activity patterns. Neuron 61:635–648.

Niell CM, Stryker MP (2008) Highly selective receptive fields in mouse visual cortex. J Neurosci 28:7520–7536.

Pallas SL (2017) The impact of ecological niche on adaptive flexibility of sensory circuitry. Front Neurosci 11:344.

Popović M, Stacy AK, Kang M, Nanu R, Oettgen CE, Wise DL, Fiser J, Van Hooser SD (2018) Development of cross-orientation suppression and size tuning and the role of experience. J Neurosci 38:2656–2670.

Prusky GT, Douglas RM (2003) Developmental plasticity of mouse visual acuity. Eur J Neurosci 17:167–173.

Rhoades RW, Chalupa LM (1978) Receptive field characteristics of superior colliculus neurons and visually guided behavior in dark-reared hamsters. J Comp Neurol 177:17–32.

Rochefort Nathalie L, Narushima M, Grienberger C, Marandi N, Hill Daniel N, Konnerth A (2011) Development of direction selectivity in mouse cortical neurons. Neuron 71:425–432.

Roy A, Christie IK, Escobar GM, Osik JJ, Popović M, Ritter NJ, Stacy AK, Wang S, Fiser J, Miller P, Van Hooser SD (2018) Does experience provide a permissive or instructive influence on the development of direction selectivity in visual cortex? Neural Devel 13(1):16.

Samonds JM, Feese BD, Lee TS, Kuhlman SJ (2017) Nonuniform surround suppression of visual responses in mouse V1. J Neurophysiol 118:3282–3292.

Sengpiel F, Stawinski P, Bonhoeffer T (1999) Influence of experience on orientation maps in cat visual cortex. Nature Neurosci 2:727–732.

Tanaka S, Miyashita M, Ribot J (2004) Roles of visual experience and intrinsic mechanism in the activity-dependent self-organization of orientation maps: theory and experiment. Neural Networks 17:1363–1375.

Van Hooser SD, Escobar GM, Maffei A, Miller P (2014) Emerging feed-forward inhibition allows the robust formation of direction selectivity in the developing ferret visual cortex. J Neurophysiol 111:2355–2373.

Wagor E, Mangini NJ, Pearlman AL (1980) Retinotopic organization of striate and extrastriate visual cortex in the mouse. J Comp Neurol 193:187–202.

Wang L, Sarnaik R, Rangarajan K, Liu X, Cang J (2010) Visual receptive field properties of neurons in the superficial superior colliculus of the mouse. J Neurosci 30:16573–16584.

White LE, Coppola DM, Fitzpatrick D (2001) The contribution of sensory experience to the maturation of orientation selectivity in ferret visual cortex. Nature 411:1049–1052.

Wiesel TN, Hubel DH (1963) Single cell responses in striate cortex of kittens deprived of vision in one eye. J Neurophysiol 26:1003–1017.

Wiesel TN, Hubel DH (1965) Comparison of the effects of unilateral and bilateral eye closure on cortical unit responses in kittens. J Neurophysiol 28:1029–1040.

Zhang M, Liu S, Wang X, Ye B, Ni Z, Ma J, Liu Y, Liu A, Zhang Z, Yu J-M, Tao W, Cao P (2026) Green light relieves stress-induced anxiety-like behaviors via visual-to-prefrontal projections in mice. Cell Reports 45.

